# CDK8 Inhibition Releases the Muscle Differentiation Block in Fusion-driven Alveolar Rhabdomyosarcoma

**DOI:** 10.1101/2025.07.14.663986

**Authors:** Susu Zhang, Kathleen Engel, Assil Fahs, Clare F. Malone, Kenneth Ross, Marissa Just, Brian Guedes, Diyana Granum, Kristianne M. Oristian, Alexander Kovach, Gabriela Alexe, Giulia Digiovanni, Leen Barbar, Rex Bentley, Christian Cerda-Smith, Ozgun Le Roux, Elizabeth Mendes, Seth P. Zimmerman, Matthew Rees, Jennifer Roth, Jack F. Shern, Kris C. Wood, Christopher M. Counter, Corinne M. Linardic, Kimberly Stegmaier

## Abstract

Alveolar rhabdomyosarcoma (aRMS) is a fusion-driven pediatric cancer with poor survival and limited therapeutic options. To uncover novel vulnerabilities, we employed complex-based analysis of the DepMap functional genomic data, identifying CDK8 as a dependency in aRMS. Both *CDK8* knockout and pharmacologic inhibition impaired tumor cell growth and induced myogenic differentiation *in vitro* and *in vivo*. Compared to genetic loss, CDK8 inhibition induced more dynamic transcriptional changes. With a genome-scale CRISPR-Cas9 drug modifier screen, we determined that the maximal anti-tumor activity of the CDK8 inhibitor requires the presence of the Mediator kinase module and transcriptional cooperation with the SAGA complex. We further identified SIX4 as a key transcription factor mediating CDK8 inhibitor-induced transcriptional activation of myogenic differentiation genes and tumor cell proliferation. These findings suggest a distinct gain-of-function mechanism of the CDK8 inhibitor and establish a strong rationale for CDK8 inhibition as a differentiation-inducing therapeutic strategy in aRMS.

**STATEMENT OF SIGNIFICANCE:** We provide a framework for uncovering therapeutic targets by network-based analysis of functional genomic screens. We identify CDK8 as a druggable target in aRMS and determine that CDK8 inhibition drives myogenic differentiation and impairs tumor progression via a collaborative mechanism involving the Mediator kinase module, SAGA complex, and SIX4.

## INTRODUCTION

Transcription factors are frequently dysregulated in human cancer leading to abnormal transcriptional programs^1–3^. Chromosomal translocations may result in the formation of transcription factor fusion oncoproteins that retain functional domains from two distinct proteins. Fusion oncoproteins are particularly common in pediatric cancers such as PAX3::FOXO1 in alveolar rhabdomyosarcoma (aRMS)^4^. In contrast to kinase-driven cancers, where small molecule inhibitors have shown success, pediatric cancers driven by fusion transcription factors present unique therapeutic challenges due to the lack of easily targetable structural pockets and intrinsically disordered domains. Additionally, while adult cancers often harbor a broad spectrum of mutations that can be therapeutically exploited, pediatric fusion-driven cancers tend to exhibit a low mutational burden, meaning fewer potential targets for precision medicine approaches^5,6^. Moreover, despite the transformative clinical success of PD-1-targeting immune checkpoint inhibitors in a subset of adult cancers, this treatment strategy has generally not benefited children with cancer^7,8^. Across four trials of monotherapies with nivolumab, pembrolizumab, atezolizumab and avelumab, 97% of the pediatric patients with solid tumor did not achieve even a partial response, and the combination of immune checkpoint inhibitors with other standard of care therapies did not improve efficacy^9–12^. Other immunotherapies, such as chimeric antigen receptor (CAR) T-cell therapies, have shown promise in hematologic malignancies, particularly B-cell malignancies, but have had only limited activity in children with solid tumors. While the expression of individual target antigens benefits subpopulations of patients, for example, GD2 in neuroblastoma or CD19 in B-cell malignancies, the lack of specificity of most surface antigens for tumor versus normal tissues has limited this strategy in the case of solid and brain tumors^13^.

As one of the most difficult to treat childhood cancers, more than 80% of the alveolar rhabdomyosarcoma (aRMS) cases harbor a PAX3/7::FOXO1 fusion, which acts as a highly active novel transcription factor with altered transcriptional properties compared to either transcription factor alone. Though more than 10,000 potential PAX3::FOXO1 regulatory sites were identified in a recent publication, only about 200 gene targets were directly transcriptionally activated by PAX3::FOXO1 based on nascent RNA sequencing, and most of these direct target genes are also transcription factors^14^. Due to the quiet mutational landscape of aRMS and the challenges in targeting transcription factors, either the fusion itself or its downstream targets, current therapy for patient with aRMS is still based on conventional strategies including chemotherapy, surgery, and radiotherapy, all of which are toxic. Efforts to improve therapeutic response in high-risk patients have included multi-agent, interval compressed therapy, high-dose chemotherapy with stem cell rescue, oral maintenance regimens, and novel targeted therapies. The overall 5-year survival, however, remains poor, and approximately one-third of the children will relapse.

Advancements in understanding dependencies in pediatric cancers have uncovered critical vulnerabilities within transcriptional machinery, offering promising therapeutic inroads. BET proteins, such as BRD4, have been recognized for their role in regulating transcriptional elongation and enhancer function through regulation of super-enhancers^15,16^. BET inhibitors, however, were met with the challenge of on-target toxicity including thrombocytopenia, and thus a narrow therapeutic window^17,18^. In contrast, menin inhibitors have emerged as a particularly notable therapeutic success. Menin serves as a key component of the MLL/COMPASS complex and plays an essential role in sustaining KMT2A-fusion transcriptional programs in acute leukemia as well as HOX/MEIS programs in *NPM1* mutant acute myeloid leukemia (AML)^19,20^. Potent menin inhibitors have demonstrated significant clinical efficacy, earning FDA-approval for *KMT2A*-fusion leukemias, marking a breakthrough in the therapeutic targeting of transcriptional co-regulators. Notably, these molecules promote myeloid differentiation in these AML cells, a feature thought to be critical to their efficacy.

Comprehensive datasets, such as Broad Institute’s Dependency Map (DepMap), have emerged as powerful tools for identifying promising single candidate therapeutic targets in cancer. Coupling the DepMap with network-based analysis allows the identification of co-dependencies, essential protein complexes and signaling nodes that are critical for cancer cell survival. This approach enables the prioritization of candidate targets based on their functional complexes and interactions within oncogenic pathways^21^. For example, the SAGA (Spt-Ada-Gcn5-acetyltrasferase) complex was reported as a dependency in *MYCN*-amplified neuroblastoma, which is driven by an aberrant oncogenic transcriptional program^22^. Several members of the complex are modest dependencies, each which individually might have been overlooked, but analysis of complexes highlighted this important neuroblastoma epigenetic regulator. This work also highlighted a framework for uncovering novel druggable targets, particularly in pediatric cancers with low mutational burdens and driven by dysregulated transcriptional machinery.

In the current study, we employed an analytical approach to identify dependencies on complexes and nominated the Mediator complex as a dependency in aRMS. We demonstrate that CDK8, a key component of the Mediator kinase module, represents a clinically actionable target in fusion-positive aRMS. Notably, inhibition of CDK8, using a small molecule inhibitor currently in clinical trials for other diseases, induced *in vivo* differentiation of aRMS tumor cells. Our findings indicate that CDK8 primarily acts as a transcriptional repressor in aRMS, directly repressing the expression of a subset of genes involved in terminal muscle differentiation. Upregulation of the transcription factor SIX4, involved in muscle differentiation, is essential to the activity of these CDK8 inhibitors. Most CDK8 inhibitors are ATP-competitive compounds that bind directly to the ATP-binding pocket to halt kinase activity^23,24^. Our findings reveal that CDK8 inhibitors elicit a more potent phenotype than CDK8 genetic deletion, and that their efficacy dependents on the presence of an intact CDK8 kinase module. These observations support a novel paradigm of kinase inhibitor trapping, in which the physical presence of the intact but enzymatically inhibited kinase target, and not just inhibition of its catalytic activity alone, is essential for the compound’s full potency. Furthermore, the efficacy of CDK8 inhibition is also dependent on the acetyltransferase activity of the SAGA complex, highlighting a cooperative interaction between these transcriptional regulators in the context of the CDK8 inhibitors.

## RESULTS

### Mediator complex catalytic component CDK8 is a selective dependency in aRMS

To identify novel pharmacologic targets in aRMS, we analyzed the Broad Institute’s Cancer Dependency Map (DepMap) dataset, which includes genome-wide CRISPR-Cas9 screens performed in over 1,000 human cancer cell line models^25^. Utilizing single-sample gene set enrichment analysis (ssGSEA), we systematically assessed protein complex and pathway dependencies in aRMS. We computed the ssGSEA scores to compare aRMS with all other cancer types in DepMap and found that in aRMS, complexes containing members of the Mediator complex [ARC (Activator-recruited cofactor), ARC-L, and ARC92-Mediator] are top hits in the ssGSEA analysis based on CORUM-defined complexes **(Fig. 1A)**. We then compared the DepMap dependency scores for the ARC-L-Mediator complex across all tested cancer cell lines, and aRMS also ranked as the disease most dependent on this complex **(Fig. S1A)**. Based on the enriched genes in the ssGSEA ARC-L/ARC92-Mediator complex data set (**Table S1A**), we then focused on components of the Mediator kinase module^26–28^. The analysis of DepMap also nominated aRMS as the disease most dependent on CDK8, the catalytic subunit of the Mediator kinase module, compared to over 1,000 tested cell lines across different types of cancers. In contrast, embryonal rhabdomyosarcoma (eRMS), the vast majority of which does not express a fusion oncoprotein, ranks like other non-CDK8 dependent malignancies **(Fig. 1B)**. In addition, when focusing on druggable dependencies, we analyzed the DepMap dependency scores of all kinases and compared fusion positive aRMS to all other tested models and found that CDK8 also scored among the top kinase hits **(Fig. 1C)**. Importantly, we found that other components of the Mediator kinase module, CCNC, MED12, and MED13, serving as essential scaffold to stabilize the kinase module within the Mediator complex, also scored as strong selective dependencies in aRMS, underscoring the critical role of this module in aRMS **(Fig. 1D)**.

**Figure 1.**
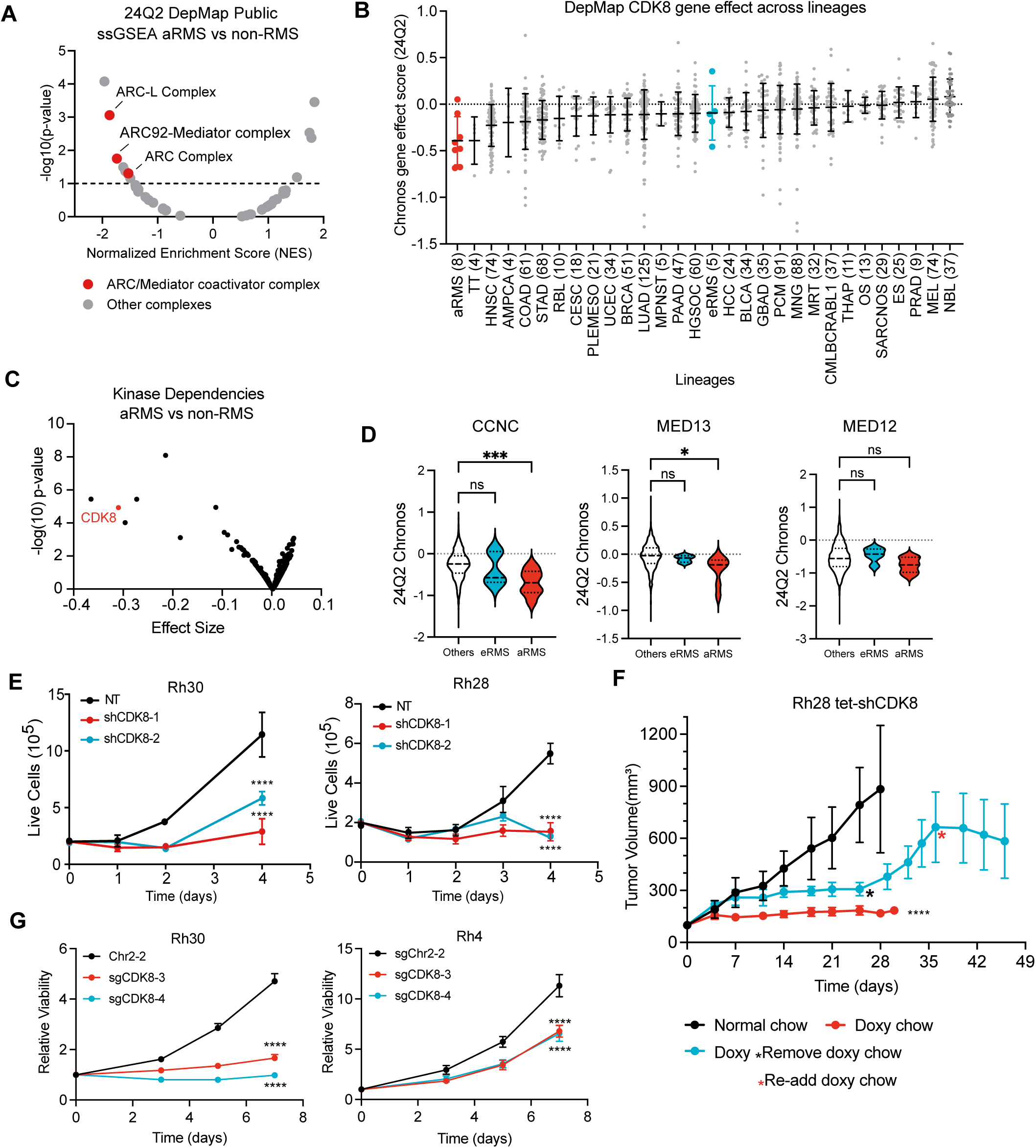
Mediator complex catalytic component CDK8 is a selective dependency in aRMS. **A.** Volcano plot of ssGSEA on genome-wide differential effect size of CORUM complexes comparing aRMS to other non-RMS tumor cell lines. Red indicates Mediator complex. **B.** Distribution of *CDK8* gene effect score across different cancer cell lines from the Broad Institute’s CRISPR Dependency Map (24Q2). **C.** Dot plot of kinase dependencies in the Broad Institute’s CRISPR Dependency Map comparing fusion-positive RMS to all other cancer cell lines. *CDK8* is highlighted in red. **D.** Violin plots showing distribution of *CCNC*, *MED13*, and *MED12* gene effect score from the Broad Institute’s CRISPR Dependency Map (24Q2) comparing the fusion-positive aRMS and fusion-negative eRMS with all other indicated cancer cell lines. aRMS is highlighted in red and eRMS is highlighted in blue. **E.** shRNA-mediated suppression of CDK8 by two different shRNAs impairs Rh30 and Rh28 aRMS cell growth *in vitro*. Cell numbers were determined by trypan blue live cell counting. Data are presented as mean ± SEM (*: *p*<=5.0e-02, **: *p*<=1.0e-02, ***: *p*<= 1.0e-03, ****: *p*<=1.0e-04). **F.** Line graph showing mean subcutaneous tumor volume (mm3) formed by Rh28 cells after treatment with inducible knock down of CDK8 using shRNA. Data are presented as mean ± SEM (*: *p*<=5.0e-02, **: *p*<=1.0e-02, ***: *p*<= 1.0e-03, ****: *p*<=1.0e-04). **G.** CRISPR-mediated knockout of *CDK8* by two different gRNAs impairs Rh30 and Rh4 aRMS cell growth *in vitro*. Relative growth was assessed by CellTiter-Glo after CRISPR knockout. Data are presented as mean ± SEM (*: *p*<=5.0e-02, **: *p*<=1.0e-02, ***: *p*<= 1.0e-03, ****: *p*<=1.0e-04).

Building on these findings, we next evaluated the RNA level of *CDK8* in different cancer cell lines using the CCLE 23Q4 dataset and in pediatric tumors using the Treehouse V11 dataset^29^. The average *CDK8* transcript levels in RMS are notably elevated compared to other cancer types **(Fig. S1B,C)**. Protein-level validation using patient-derived xenograft (PDX) tumor microarray samples demonstrated robust expression of CDK8 in aRMS PDX tumors (**Fig. S1D**). In addition, PAX3::FOXO1-expressing aRMS cell lines exhibited significantly higher CDK8 protein levels compared to non-transformed primary human skeletal muscle cells (**Fig. S1E**).

### CDK8 inhibition slows aRMS cell growth *in vitro* and *in vivo*

To evaluate the functional consequences of CDK8 loss of function in aRMS, we employed both genetic knockdown and pharmacological approaches. Two independent shRNAs targeting *CDK8* were used to perturb *CDK8* expression genetically. CDK8 was down regulated effectively at the protein level in Rh28 and Rh30, two fusion positive aRMS cell lines (**Fig. S1F**). Genetic knockdown of *CDK8* resulted in a marked depression of cell viability (**Fig. 1E**). More importantly, using a shRNA conditional knockdown system in the Rh28 cell line, we confirmed that *CDK8* depletion significantly inhibited tumor cell growth *in vivo* (**Fig. 1F, S1G**). Similarly, CRISPR-mediated genetic knockout of *CDK8* led to substantial growth inhibition and down-regulation of its kinase target pSTAT1^S727^ phosphorylation^30^ compared to the cells transduced with control guide RNAs in aRMS cell lines, Rh30 and Rh4 (**Fig. 1G, S1I**). We noticed some variation of growth inhibition among different cell lines, such as RHJT, which demonstrated a more modest response to *CDK8* depletion (**Fig. S1H, I**).

To evaluate the relationship between genetic dependency on *CDK8* and pharmacological inhibition, we compared DepMap gene-effect scores for *CDK8* knockout with PRISM (Profiling Relative Inhibition Simultaneously Mixtures) log_2_ fold-change data for the CDK8 inhibitor BI-1347. In aRMS, we observed a strong positive correlation (R=0.68) between these two metrics, whereas the correlation was lower in embryonal RMS (R=0.32) and other cancer cell lines (R=0.21) (**Fig. 2A**). Therefore, in addition to genetic manipulation of CDK8 expression, we tested three selective CDK8 kinase inhibitory small molecules, including BI-1347, SEL-120-34A, and JH-XII-178, all of which demonstrated on-target activity by reducing STAT1^S727^ phosphorylation (**Fig. 2B**). In the tested aRMS cell lines, all three CDK8 inhibitors impaired cell growth, while the inactive analog of BI-1347, known as BI-1374, exhibited no effect (**Fig. 2C, D**). Following our observation of the cell growth defect, we assessed for apoptosis by examining Caspase-Glo expression, cleaved PARP levels, Annexin V staining, and cell cycle phase. Although we observed a mild increase in apoptosis and G2M-phase cell cycle arrest after genetic or pharmacological inhibition of CDK8, these effects were insufficient to fully explain the observed growth inhibition (**Fig. S1J-M**). To further characterize the impact of CDK8 inhibition, we utilized Incucyte live-cell imaging to monitor cell proliferation. Compared to DMSO and an inactive analog, long-term exposure to the CDK8 inhibitor BI-1347 slowed aRMS cell proliferation, suggesting that the primary mechanism of CKD8 inhibition-mediated growth suppression involves reduced proliferative capacity rather than apoptosis or cell cycle arrest (**Fig. 2E, F**). Last, we asked whether CDK8 kinase inhibition would confer clinical benefits for aRMS *in vivo*. We therefore tested the CDK8 inhibitor SEL-120-34A, which has been granted orphan drug designation by the FDA for the treatment of AML and high-risk MDS, in an Rh30 cell line xenograft model^31^. We observed that treatment with SEL-120-34A decreased tumor growth *in vivo* without affecting animal weight (**Fig. 2G, S1N**). Consistent with *in vitro* observations, tumors from SEL-120-34A-treated mice exhibited a mild increase in cleaved caspase 3 staining (**Fig. S1O**). These data suggest that CDK8 inhibition impairs tumor progression *in vivo* as well as *in vitro*.

**Figure 2.**
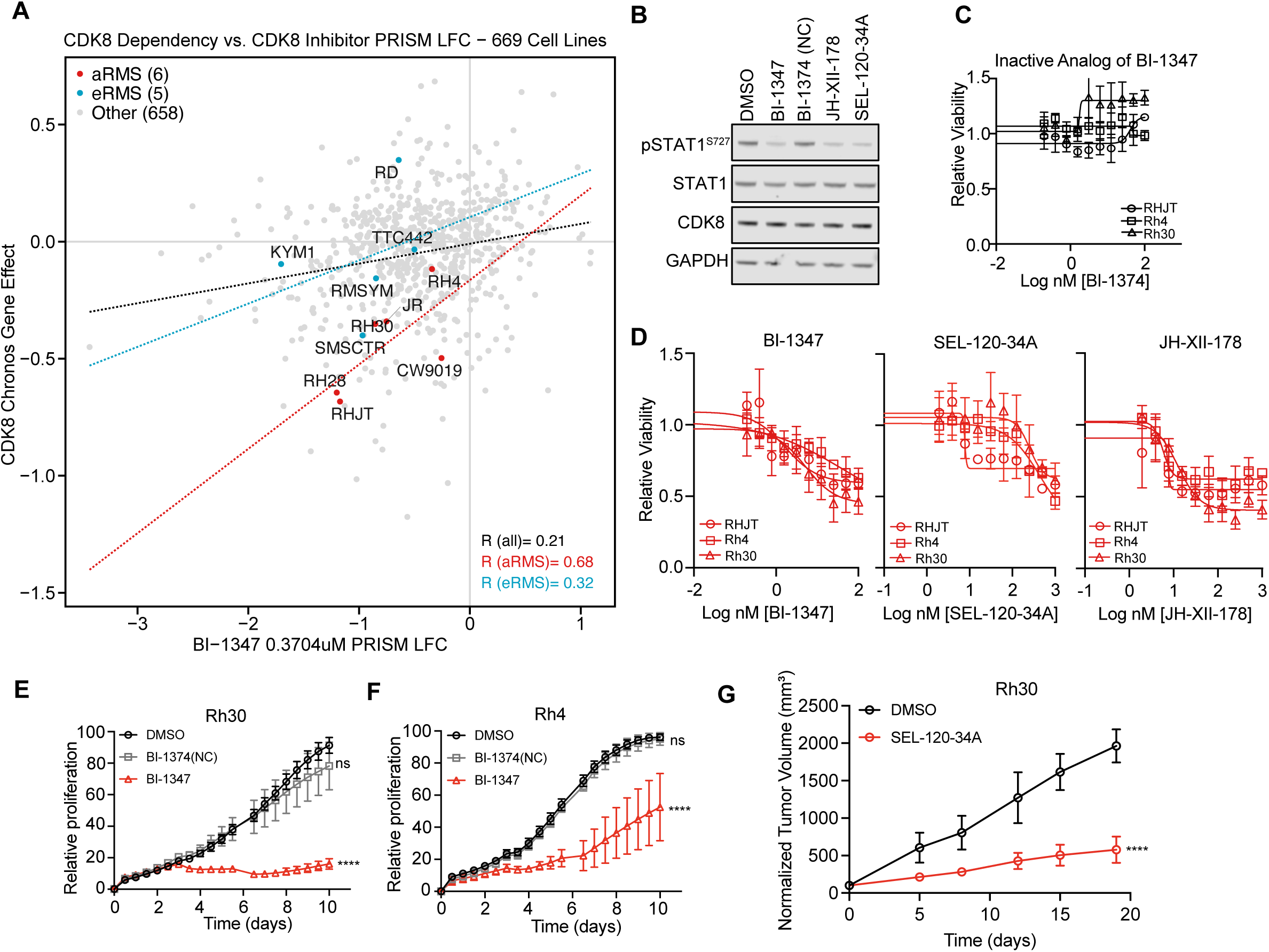
Pharmacologic inhibition of CDK8 in fusion-positive aRMS impairs cell growth *in vitro* and *in vivo*. **A.** Correlation between *CDK8* gene dependency and sensitivity to the CDK8 inhibitor BI-1347 in 669 cancer cell lines (y-axis showing CDK8 dependency score; x-axis showing PRISM LFC value of BI-1347 treatment). Linear regression lines and Pearson correlation coefficients (R) are shown for all cell lines (black), aRMS (red), and eRMS (blue). **B.** Western blot analysis showing CDK8 inhibition by small molecules as assessed by STAT1 phosphorylation at serine 727. **C, D.** Dose response curves of inactive BI-1347 analog (BI-1374) (C) and three pharmacologic CDK8 inhibitors (D), BI-1347, SEL-120-34A, and JH-XII-178, at day 7 of treatment in Rh30, Rh4 and RHJT cell lines. **E, F.** Live cell proliferation assessed by Incucyte for Rh30 (E) and Rh4 (F) cells after treatment with vehicle DMSO (black), the CDK8 inhibitor BI-1347 (red) and its inactive analog BI-1374 (gray). Data were normalized to DMSO. Data represent means ± SEM (n=6, *: *p*<=5.0e-02, **: *p*<=1.0e-02, ***: *p*<= 1.0e-03, ****: *p*<=1.0e-04). **G.** Line graph reveals mean subcutaneous tumor volume (mm^3^) formed by Rh30 cells after treatment with the CDK8 inhibitor SEL-120-34A. Data are presented as mean ± SEM (*: *p*<=5.0e-02, **: *p*<=1.0e-02, ***: *p*<= 1.0e-03, ****: *p*<=1.0e-04).

### CDK8 inhibition induces transcriptional activation of myogenic differentiation-associated gene targets

As CDK8 serves as the catalytic component of the Mediator kinase module, we next performed Precision nuclear Run-On transcription sequencing (PRO-seq) to determine the direct effect of CDK8 inhibition on early events in transcription, particularly RNA polymerase II (Pol II) pausing and elongation^32–34^. We analyzed RNA polymerase II occupancy using normalized changes in the gene body and by performing differential analysis. We did not observe a global change in RNA polymerase II at gene bodies between 4 to 72 hours following BI-1347 treatment in Rh30 cells, indicating that CDK8 kinase inhibition does not lead to a widespread alteration in Pol II transcriptional activity (**Fig. 3A, Fig. S2A-B**). There were very few changes noted at the 4-hour timepoint. After 24 hours of CDK8 inhibition by BI-1347, however, there were 523 genes exhibiting upregulated gene body transcription and 59 genes exhibiting downregulated gene body transcription (**Fig. 3A, Table S1B**). A similar transcriptional trend was observed at the 72-hour timepoint of BI-1347 treatment, with 575 genes exhibiting upregulated gene body transcription and 193 genes exhibiting downregulated gene body transcription (**Fig. S2B, Table S1C**). These changes at the two treatment time points were relatively consistent; the dominant effect of CDK8 inhibition was activation of transcription (**Fig. 3A, S2A-C**). When comparing differentially expressed genes identified by PRO-seq with CDK8-bound genes, we found 65% of upregulated and 44% of downregulated genes were directly associated with CDK8 binding, suggesting a strong correlation between CDK8 occupancy and its transcriptional regulatory effects (**Fig. S2D**). The heatmaps of the normalized RNA polymerase II occupancy and pausing index indicated increased occupancy along the gene body of RNA polymerase II, further suggesting increased transcriptional elongation at gene bodies (**Fig. 3C, S2E**). The steady-state RNA level as assessed by whole transcriptome RNA-seq correlated with the changes observed in the nascent transcription analysis, demonstrating concordance between nascent and steady-state RNA levels (**Fig. 3D**).

**Figure 3.**
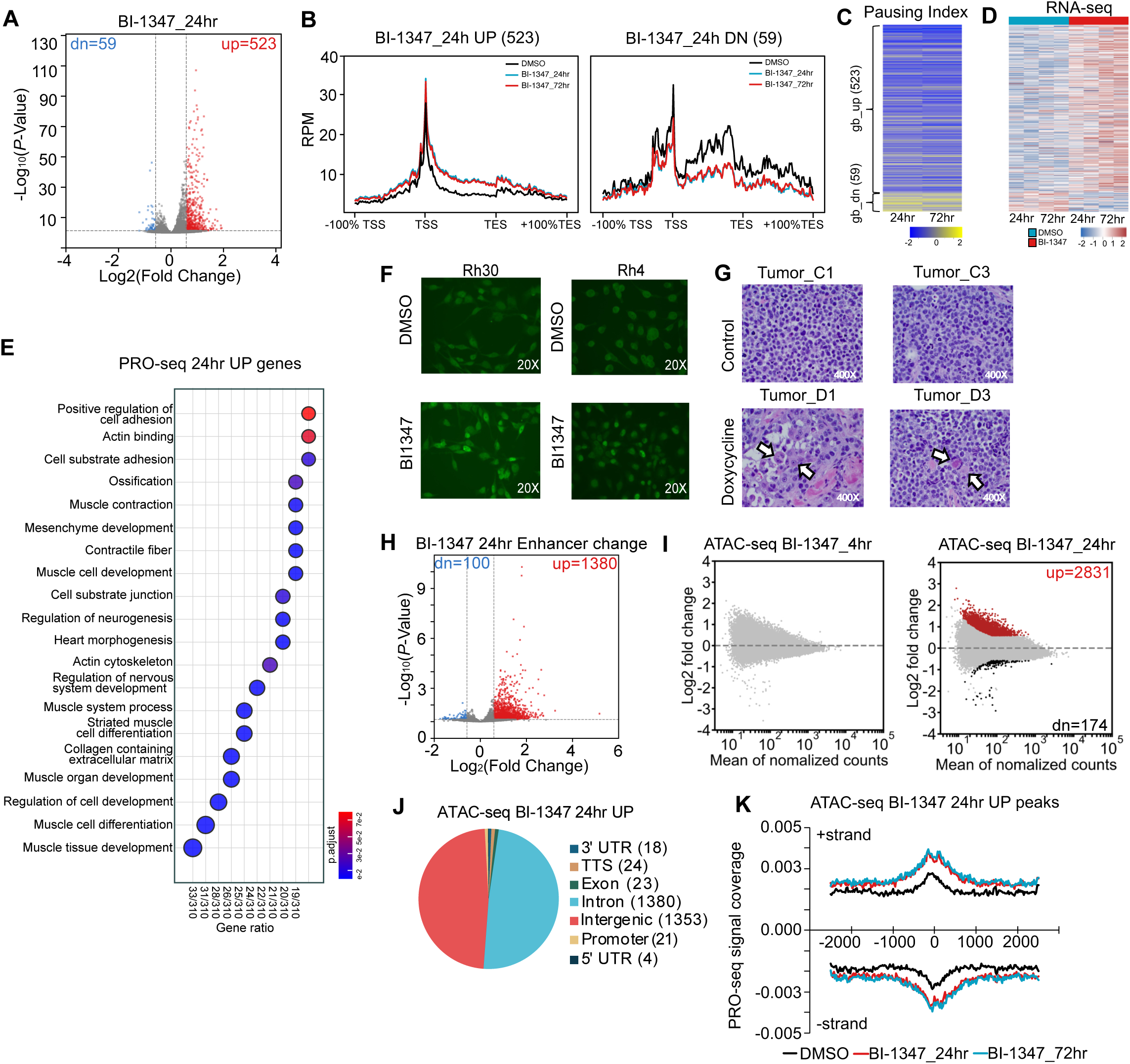
CDK8 inhibition induces transcriptional activation of myogenic differentiation-associated genes. **A.** Volcano plots showing the number of gene body changes after 24 hrs of CDK8 inhibitor BI-1347 treatment. Significantly up-regulated genes are highlighted in red; significantly downregulated genes are highlighted in blue (padj<0.05, fold change>1.5 or <-1.5). **B.** Metagene plots of PRO-seq reads of genes with significant upregulation or downregulation of gene body transcription at 24 hrs of BI-1347 treatment. **C.** Heatmap of Log_2_ transformed fold change values of PRO-seq pausing indices of genes with changes in gene body transcription at the 24 hr time point. **D.** Heatmap showing log_2_ transformed fold change of RNA-seq read counts for indicated gene sets after DMSO and BI-1347 (10 nM) treatment for 24 or 72 hrs. **E.** Bubble dot plot shows Gene Set Enrichment Analysis (GSEA) of PRO-seq gene hits that are upregulated at 24 hrs. Dot size indicates number of genes in each gene set, and dot color indicates *p*-adjusted value. **F.** Immunofluorescence analysis of myogenin (green; 20X) in Rh30 (left) and Rh4 (right) cells after 7 days of DMSO or BI-1347 treatment. **G.** Representative images of H&E stain of xenograft tumors. Arrows indicate myofibrils. **H.** Volcano plots reveal the number of changes in enhancer RNA (eRNA) transcripts after 24 hrs of BI-1347 treatment. Significantly upregulated eRNAs are highlighted in red; significantly downregulated eRNAs are highlighted in blue (padj<0.05, fold change>1.5 or <-1.5) **I.** MA plots showing the changes of chromatin accessibility by ATAC-seq following 4 and 24 hrs of BI-1347 treatment. Significantly increased ATAC-seq peaks are highlighted in red; significantly decreased ATAC-seq peaks are highlighted in black (padj<0.05, fold change>1.5 or <-1.5). **J.** Pie chart showing the peak annotation of up-regulated ATAC-seq peaks to gene features at 24 hrs of BI-1347 treatment. **K.** Histogram of PRO-seq reads around up-regulated ATAC-seq peaks at indicated time points of BI-1347 treatment.

Gene Set Enrichment Analysis (GSEA) revealed a strong overlap between transcriptional changes driven by *CDK8* genetic knockout (sgCDK8) and BI-1347 treatment (**Fig. S2F**). To further annotate the changes induced by BI-1347, we performed GSEA on genes identified to be up regulated in the 24-hour PRO-seq data. Notably, early transcriptional activation at 24 hours of BI-1347 treatment was enriched for muscle-related pathways, including muscle tissue development, muscle cell differentiation, as well as other annotations related to muscle cell development, as identified by GSEA with C5 ontology gene sets (**Fig. 3E**). Similarly, RNA-seq data generated following *CDK8* genetic knockout highlighted genes associated with myogenesis as the top upregulated signature (**Fig. S2G**). After BI-1347 treatment, we detected noticeable increases in expression of the muscle differentiation marker myogenin as assessed by immunofluorescence (**Fig. 3F**). Importantly, upon *CDK8* genetic inhibition, we observed increased cytoplasmic eosin staining with eccentrically located nuclei structures in the xenograft tumor samples, indicative of myofibril generation, a morphological hallmark of myogenic differentiation *in vivo*^35^ (**Fig. 3G, S2H**).

While global RNA Pol II occupancy at gene bodies remained largely unchanged following BI-1347 treatment, transcriptional regulation can also occur at the enhancer level. To further elucidate the transcriptional mechanisms underlying CDK8 inhibition, we utilized the previously obtained PRO-seq data to quantify enhancer RNA (eRNA) expression following CDK8 inhibition. After 24 hours of BI-1347 treatment, there were 1,380 eRNAs with increased expression, while only 100 had decreased expression (**Fig. 3H**). We then evaluated for activated enhancers by assessing for chromatin accessible regions using ATAC-seq. Consistent with the PRO-seq analysis, while we did not observe global changes in chromatin accessibility, we did identify focal gains in chromatin accessibility (2,831 increased peaks versus 174 decreased peaks), predominantly (>90%) in intergenic or intronic regions (**Fig. 3I, 3J, S2I**). To confirm that these regions with increased accessibility were undergoing nascent transcription, we then plotted the PRO-seq signal ± 2.5kb at these upregulated ATAC-seq peaks, with the histogram showing an increase in active polymerase around these upregulated ATAC-seq regions after 24 and 72 hours of CDK8 inhibition (**Fig. 3K**). These findings suggest CDK8 inhibition does not globally impact transcriptional activities at gene bodies or enhancers, but instead selectively activates the transcription of specific genes by enhancing eRNA production and activity.

To assess whether CDK8 inhibition affects the expression or the binding to the targets of PAX3::FOXO1, the most common fusion oncoprotein driving aRMS, we first examined whether CDK8 inhibition impacts mRNA levels of the fusion gene. At a transcriptional level, we did not observe changes in *PAX3::FOXO1* gene body transcription following BI-1347 treatment for up to 72 hours of treatment (**Fig. 4A**). In addition to its role in Mediator function, CDK8 has also been reported to phosphorylate transcription factors^30^. A kinase prediction tool indicated that a phosphorylation site on wildtype FOXO1 is a predicted site of CDK8, and this site is preserved in the PAX3::FOXO1 protein (FOXO1^Ser479^) (https://gps.biocuckoo.cn/index.php). While there is currently no available antibody to interrogate this Ser479 phosphorylation site, given that the phosphorylation event might impact PAX3::FOXO1 stabilization and/or cytoplasmic/nuclear transport, we examined the levels of PAX3::FOXO1 protein after treatment with the CDK8 inhibitor BI-1347. Neither total PAX3::FOXO1 protein levels nor its nuclear or cytoplasmic distribution was notably altered by CDK8 inhibition (**Fig. S3A**).

**Figure 4.**
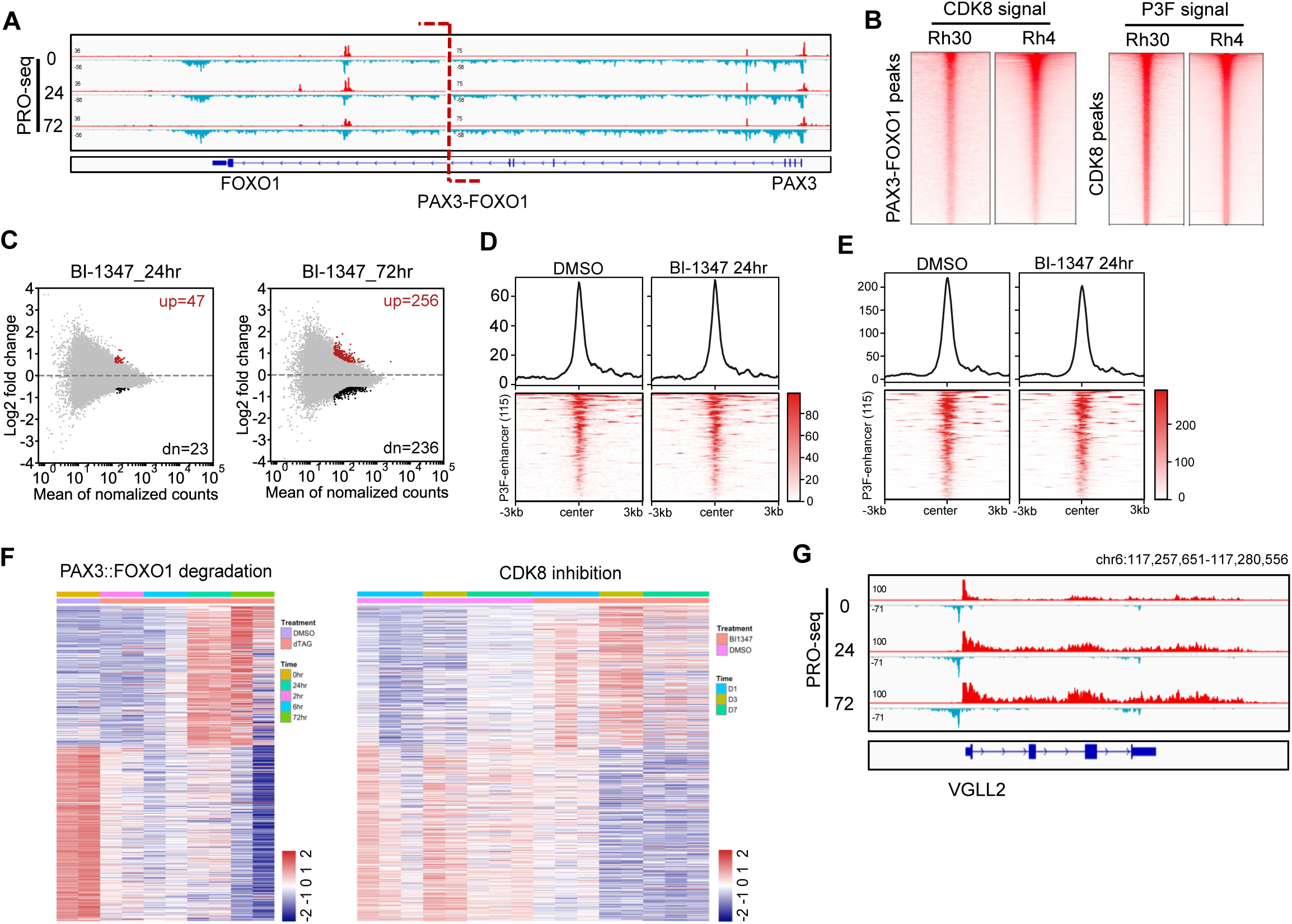
CDK8 inhibition induces a differentiation program without altering PAX3::FOXO1. **A.** IGV gene tracks showing a time course analysis of BI-1347 treatment by PRO-seq signal at the *PAX3::FOXO1* locus. B. Heatmaps of CUT&RUN analysis of CDK8 and PAX3::FOXO1 binding in Rh30 and Rh4 cell lines. C. MA plots showing changes of PAX3::FOXO1 binding following BI-1347 treatment at 24 and 72 hrs. Significantly increased PAX3::FOXO1 peaks are highlighted in red; significantly decreased PAX3::FOXO1 peaks are highlighted in black (padj<0.05, fold change>1.5 or <-1.5). **D, E.** Heatmaps of CDK8 (D) and PAX3::FOXO1 (E) signal around PAX3::FOXO1-regulated enhancers before and after 24 hrs of BI-1347 treatment. **F.** Heatmaps of log_2_ transformed fold change of RNA-seq read counts plotted by the shared gene between 24 hrs of PAX3::FOXO1 degradation and 72 hrs of CDK8 inhibition at the indicated time points. **G.** IGV gene tracks showing a time course analysis for PRO-seq signal at the *VGLL2* locus after BI-1347 treatment.

Previous studies reported CDK8 among the proteins in close proximity to endogenous PAX3::FOXO1, and CDK8 was enriched in binding at high confidence PAX3::FOXO1 regulated enhancers^14^. To verify this interaction, we first performed BioID protein-proximity labeling in HEK293T cells expressing a Myc-BirA_PAX3::FOXO1 fusion protein, as well as with mock control cells and a Myc-BirA control (**Fig. S3B**). Stable expression of the Myc-BirA_PAX3::FOXO1 fusion, the Myc-BirA, and biotinylation, were further confirmed by confocal immunofluorescent imaging (**Fig. S3C**). Streptavidin affinity pulldown followed by quantitative LC/MS/MS identified CDK8 peptides as one of the top hits other than the PAX3::FOXO1 peptides (**Fig. S3D**), consistent with the previously reported PAX3::FOXO1 binding proteins, including CDK8, identified by APEX2 proximal labeling in Rh30 cells^14^. Moreover, given that the Mediator complex is involved in facilitating promoter-enhancer interactions, and CDK8 showed enrichment at PAX3::FOXO1 regulated enhancers, we next evaluated whether PAX3::FOXO1 binding changes globally or at PAX3::FOXO1 regulated enhancers after CDK8 inhibition or knockout. Consistent with the previously reported study of PAX3::FOXO1 endogenous degradation, CDK8 co-localized with PAX3::FOXO1^14^ (**Fig. 4B**). To more accurately detect PAX3::FOXO1 changes on chromatin, we next used an epitope-tagged PAX3::FOXO1_FKBP-HA cell line and HA antibody to map PAX3::FOXO1 genome binding following CDK8 inhibition^14^. We found only minimal changes in PAX3::FOXO1 binding after CDK8 inhibition at 24 and 72 hours (**Fig. 4C**). For the high-confidence PAX3::FOXO1-regulated enhancers defined previously^14^ (115 peaks), neither CDK8 binding (**Fig. 4D**) nor PAX3::FOXO1 (**Fig. 4E**) binding was affected by CDK8 inhibition. These results indicate that the impact of CDK8 inhibition on FP-RMS cell biology may not function directly through PAX3::FOXO1-regulated core enhancers. However, RNA-seq revealed shared transcriptional changes between CDK8 inhibition and PAX3::FOXO1 degradation over long-term treatment, manifested by an enrichment in genes associated with muscle differentiation, such as *VGLL2* (**Fig. 4F, G**).

### Interruption of the SAGA complex is sufficient to rescue the impaired growth induced by CDK8 inhibitors

To gain insight into mechanisms of resistance or sensitization to CDK8 inhibitors in aRMS, we performed a genome-scale CRISPR-Cas9 drug modifier screen against the CDK8 inhibitor BI-1347. We generated an Rh30-Cas9-mCherry cell line and infected it with the Avana-4 sgRNA library^36^. The transduced cells were further divided into two groups, DMSO and BI-1347, and treated for 21 days (**Fig. S4A**). We collected cells on both days 14 and 21 to measure sgRNA abundance and identify potential modifiers. We first intersected the top 10 hits detected from days 14 and 21 to confirm the robustness and reproducibility of the screen (**Fig. 5A, S4B,C, and Tables S1D,E**). We further analyzed the pathways enriched with resistance to 21 days of BI-1347 treatment using the GSEA C5 gene sets. The most enriched gene signatures were associated with SAGA (Spt-Ada-Gcn5 acetyltransferase) coactivator complex or transcriptional activation (**Fig. 5B**). Notably, three of the top six shared genes are components of the SAGA coactivator complex: TAF5L belonging to the core module, and TADA2B and KAT2A forming part of the histone acetyltransferase (HAT) module (**Fig. 5C**). Additionally, SGF29 from the HAT module and SUPT20H from the core module were also among the shared hits identified at both days 14 and 21 (**Fig. 5A, 5C, S4B, S4C**). Resistance hits (genes whose knockout rescued growth under BI-1347 treatment) scored higher than sensitizing hits (genes whose knockout enhanced BI-1347 sensitivity) in our screen (**Fig. 5A, S4B, S4C**). To validate these findings, we conducted cell proliferation assays using Incucyte live-cell imaging to evaluate BI-1347 treatment in the context of knockout of members of the SAGA complex. We examined three CDK8 inhibitors: BI-1347, SEL-120-34A and JH-XII-178. As shown previously in Fig. 2D, all three CDK8 inhibitors slowed proliferation. Then, we used two guide RNAs to target *TAF5L* and *TADA2B*, and a KAT2A/B Proteolysis Targeting Chimera (PROTAC), GSK699, to degrade KAT2A and KAT2B^37^. Consistent with the screen results, all three SAGA components when depleted rescued the proliferation defect induced by CDK8 inhibitors, consistent with the screen results (**Fig. 5D-5F, S4D-S4I**). Furthermore, CDK8 inhibition-induced transcriptional activation of muscle differentiation-associated genes, including *RUNX1*, *SEMA3D* and *VGLL2*, was reversed upon disruption of TAF5L, TADA2B, or KAT2A/B (**Fig. 5G, S4J, S4K**). Together with the PRO-seq data, these findings support a transcriptional repressive role of CDK8 in aRMS and the requirement of SAGA complex for induction of differentiation associated genes upon CDK8 inhibition in aRMS.

**Figure 5.**
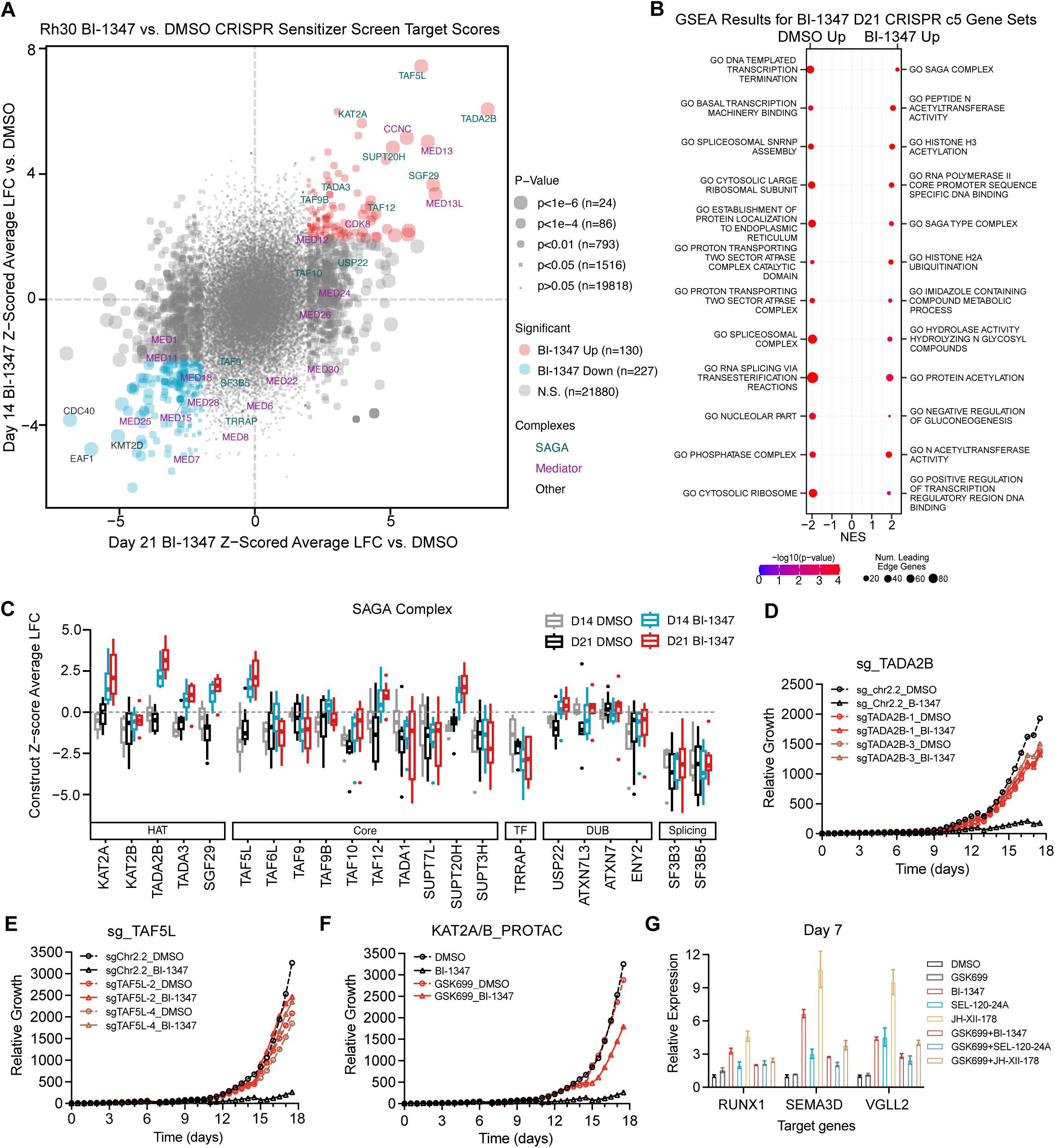
Interruption of the SAGA complex rescued the inhibitory effect of CDK8 inhibitors on growth of aRMS cells. **A.** Scatter plot showing the Z-scored average log2 fold change (LFC) of gene knockout effects in Rh30 cells treated with BI-1347 versus DMSO at day 14 (y-axis) and day 21 (x-axis) from genome-wide CRISPR-Cas9 screens. Each point represents an individual gene; dot size corresponds to statistical significance, and dot color indicates classification. **B.** Bubble dot plot of GSEA for gene hits scoring at day 21 in the CRISPR-Cas9 BI-1347 drug modifier screen ranked by C5 gene sets. **C.** Box plots showing construct-level Z-score averages for individual genes in the SAGA complex from the genome-wide CRISPR-Cas9 screen in Rh30 cells treated with DMSO (gray/black) or BI-1347 (blue/red) for 14 days (gray and blue) or 21 days (black and red). Genes are grouped by SAGA functional modules. **D-F.** Live cell proliferation assessed by Incucyte for BI-1347+/-sgTADA2B (**D**), BI-1347+/-sgTAF5L (**E**), and BI-1347+/-GSK699 (**F**). **G.** Quantitative real-time TaqMan qPCR analysis of *RUNX1*, *SEMA3D*, and *VGLL*2 expression at day 7 following treatment with vehicle control (DMSO), CDK8 inhibitors, or GSK699, and the indicated combination treatments. Expression levels were normalized to *GAPDH* gene expression and shown relative to DMSO control. Data represent means ± SEM (n=6).

### CDK8 inhibition induces increased binding of SIX4 and the SAGA complex to terminal muscle differentiation genes

Our findings indicate that CDK8 inhibition predominantly leads to transcriptional activation, with the SAGA transcriptional coactivator complex playing a critical role in mediating this effect. We next asked whether there is a specific transcription factor(s) involved in this regulation. Our screen on both days 14 and 21 identified the transcription factor SIX Homeobox 4 (SIX4), reported to play a critical role in myogenic differentiation, as a top transcription factor gene hit^38,39^ (**Fig. S4C, Table S1A, S1B**). Incucyte analysis showed that knockout of *SIX4* partially rescued the growth inhibition caused by BI-1347 (**Figs. 6A, S5A**). Total protein levels of SIX4 increased after treatment with each of the three CDK8 inhibitors (**Fig. 6B**), and SIX4 was predominantly localized to the nucleus (**Fig. 6C, 6D**). Consistently, CDK8 inhibitors also elevated *SIX4* mRNA levels as detected by both PRO-seq (24 hrs LFC= 0.4) and RNA-seq (24 hrs LFC=0.687, 72 hrs, LFC=0.642, day 7=0.488) (**Fig. S5B**). We next examined the localization of SIX4 and the SAGA component TADA2B on chromatin by CUT&RUN to further characterize their roles in CDK8-mediated transcriptional regulation. Following BI-1347-mediated CDK8 inhibition, we observed a significant increase in SIX4 and TADA2B binding on chromatin (**Fig. 6E, 6F**). Peak annotation using Homer analysis revealed that nearly 90% of the increased binding peaks for both SIX4 and TADA2B were annotated to intronic or intergenic regions, while the overall SIX4 and TADA2B binding sites are evenly distributed among promoter, intron, and intergenic regions (**Fig. 6G, 6H**, **Fig. S5C, S5D).** Accordingly, we found that the increase in SIX4 and TADA2B binding with CDK8 inhibitor treatment correlated with the enhanced chromatin accessibility identified by ATAC-seq, as well as increased H3K27ac (**Fig. 6I, S5E, S5F**). Examination of key skeletal muscle development-related genes, such as *RUNX1* and *VGLL2,* illustrates the dynamics of these changes (**Fig. 6J, S5G**). By 24 hours of CDK8 inhibition, enhancer RNA (eRNA) transcription increased at upstream intergenic enhancers of *RUNX1* and upstream intergenic enhancers of *VGLL2*. These regions also exhibited upregulated binding of SIX4 and TADA2B, correlating with active transcription of *RUNX1* and *VGLL2* gene bodies (**Fig. 6J, S5G**).

**Figure 6.**
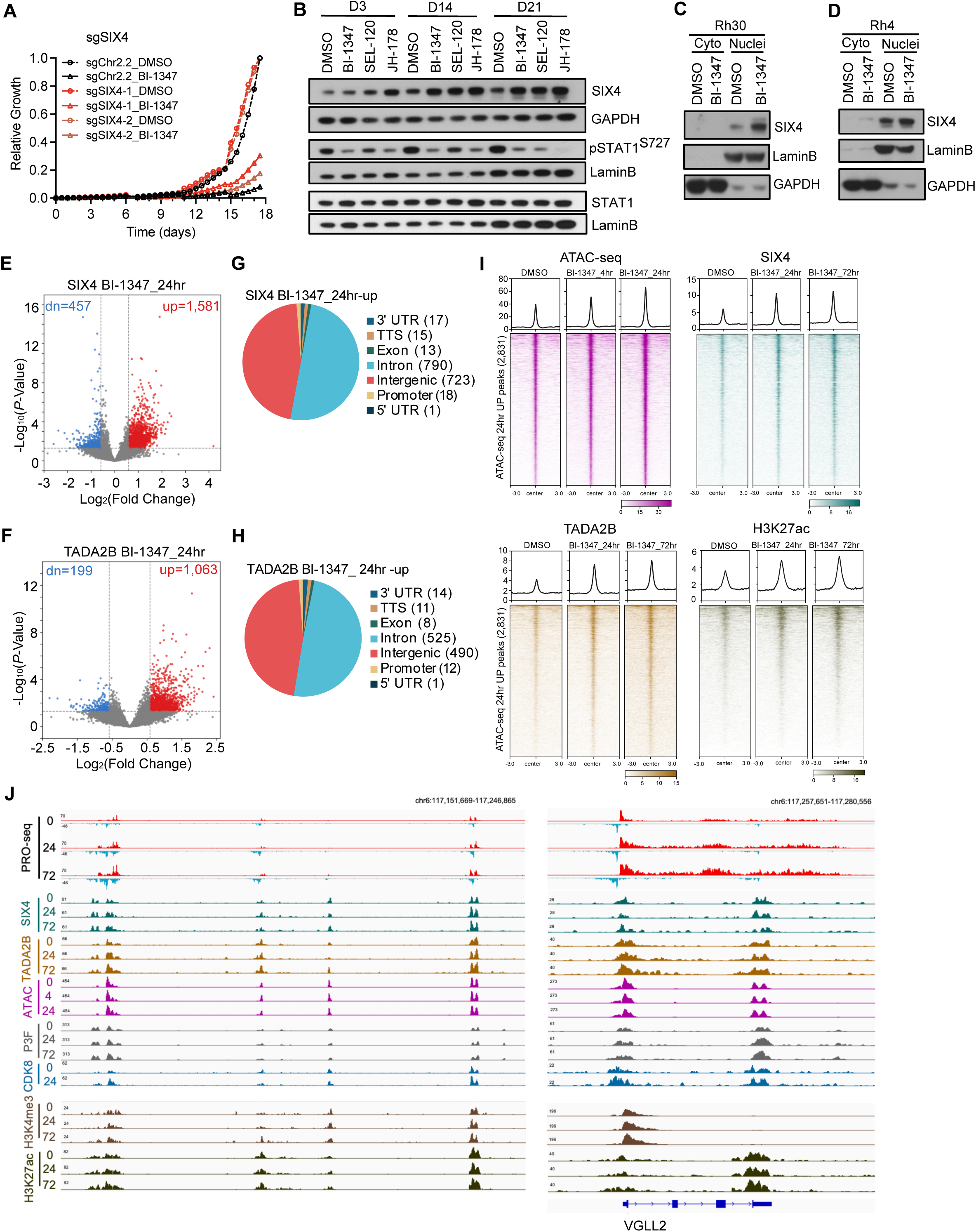
CDK8 inhibitor BI-1347 increases SIX4-driven gene expression involved in terminal muscle differentiation. **A.** Live cell proliferation assay by Incucyte for BI-1347 treatment combined with CRISPR knockout of *SIX4* with two different guide RNAs. **B.** Western blot analysis of SIX4 protein level after treatment with three CDK8 inhibitors at indicated time points. Lamin B included as a loading control. **C, D.** Western blot analysis of SIX4 protein levels from cytoplasmic and nuclear fractions from Rh30 (**C**) and Rh4 (**D**) cells treated with DMSO or BI-1347 for 14 days. GAPDH and Lamin B served as nuclear and cytoplasmic loading controls, respectively. **E, F.** Volcano showing log_2_ fold change of SIX4 (**E**) and TADA2B (**F**) genome binding sites determined by CUT&RUN analysis. Red highlights significantly up regulated sites and blue highlights significantly down regulated sites (log_2_FC 1.5, padj<0.05). **G, H.** Pie charts showing the annotation of upregulated CUT&RUN peaks for SIX4 (**G**) and TADA2B (**H**) sites after 24 hrs of BI-1347 treatment in Rh30 cells. **I.** Heatmaps of time course analysis of ATAC-seq signal, SIX4 signal, TADA2B signal, H3K27ac signal around upregulated ATAC-seq peaks at 24 hrs of BI-1347 treatment. **J.** IGV gene tracks showing the PRO-seq, SIX4, TADA2B, ATAC-seq, PAX3::FOXO1, CDK8, H3K4me3, and H3K27ac at the *VGLL2* gene body and enhancers loci at indicated time points of BI-1347 treatment.

### The CDK8 kinase module is required for CDK8 inhibitor activity

Unexpectedly, MED13 and CCNC, key components of the Mediator CDK8 kinase module, emerged as two of the top six shared rescue gene hits between days 14 and 21 in the genome-scale CRISPR-Cas9 BI-1347 drug modifier screen (**Fig. 5A, S4B, S4C**). A detailed analysis of the gRNA abundance targeting all kinase module members revealed enrichment after 21 days of BI-1347 treatment, especially for guides directed against *CCNC*, *MED13*, and *MED13L* (**Fig. 7A**). In contrast to our results for the SAGA complex and SIX4, where knockout of these genes does not affect growth in DMSO treated samples, knockout of *CDK8*, *CCNC*, *MED13*, or *MED12* impairs growth in the vehicle treated condition (**Fig. 7A**, comparing gray/black box to gray dash line), recapitulating the DepMap findings and our validation studies of the CDK8 dependency in aRMS (**Fig. 1A-G**). We validated the dependency on *CCNC* in addition to *CDK8* when comparing the abundance of gRNAs on days 0 and 21 of DMSO treatment (**Fig. 7A**, comparing gray/black box to gray dash line). In addition, Incucyte assays further substantiated the dependency on *CCNC* in the Rh30 cell line (**Figs. 7B, S6A, S6B**, comparing black dash line to two blue dish lines). When comparing the relative proliferation rate, however, genetic knockout of *CCNC* partially rescued the inhibitory effects of BI-1347 (**Fig. 7B**). RNA-seq analysis show only minor changes in the RNA levels of *CDK8, MED12, MED13, MED13L*, and *CCNC* after CDK8 inhibition or genetic knock out (**Fig. S6C**). However, the protein levels of these genes were increased after prolonged (14 days) CDK8 inhibition with three different CDK8 inhibitors (**Fig. S6D**).

**Figure 7.**
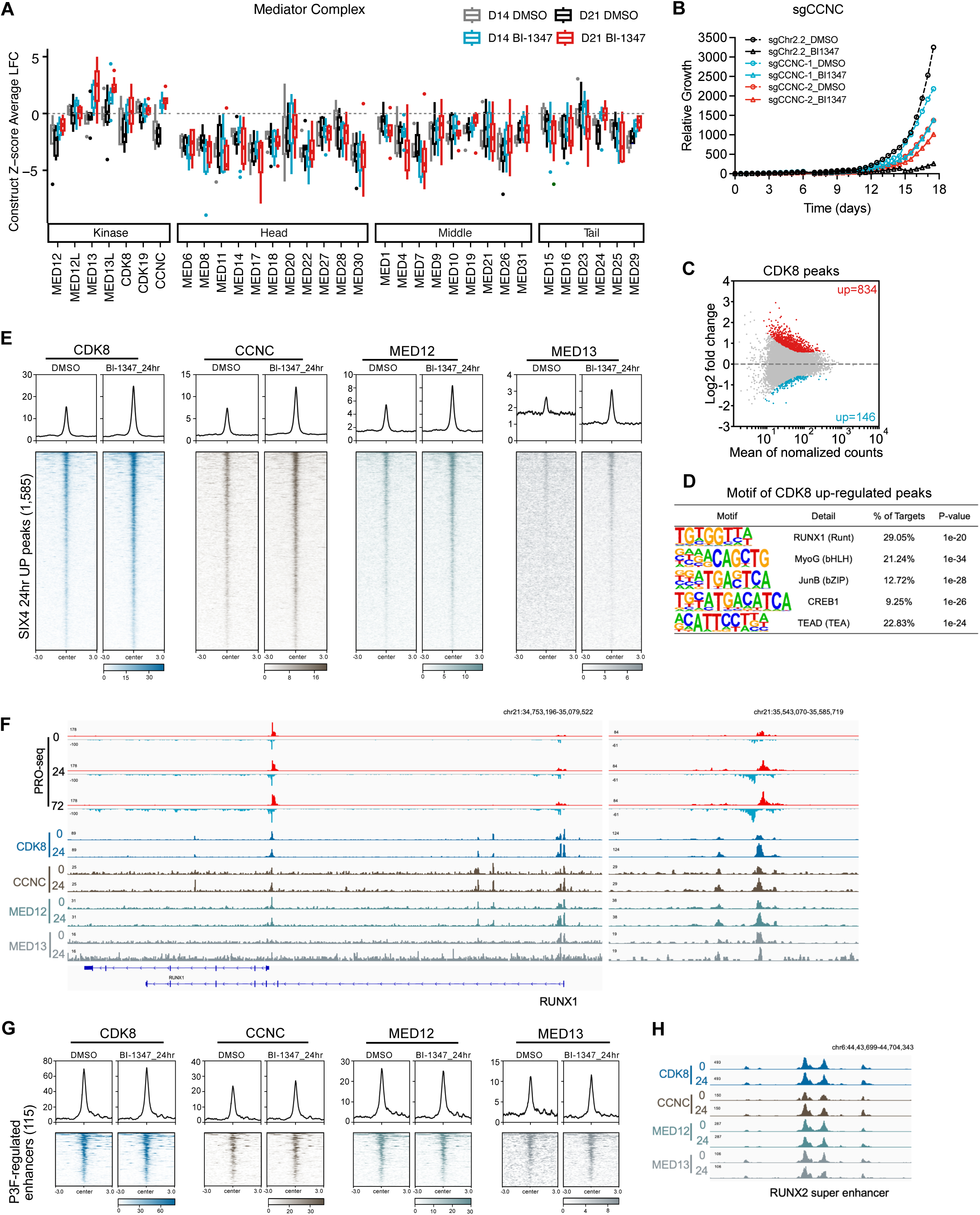
The Mediator kinase module is required for maximal CDK8 inhibitor activity. **A.** Box plots showing construct-level Z-score averages for individual genes in the Mediator complex from a genome-wide CRISPR-Cas9 screen in Rh30 cells treated with DMSO (gray/black) or BI-1347 (blue/red) for 14 days (gray and blue) or 21 days (black and red). Genes are grouped by Mediator functional modules. **B.** Live cell proliferation assessed by Incucyte for BI-1347+/-sgCDK8 (red) and BI-1347+/-sgCCNC (blue). **C.** MA plot showing changes of CDK8 binding site assessed by CUT&RUN after 24 hrs of BI-1347 treatment. Significantly increased CDK8 peaks are highlighted in red; significantly decreased CDK8 peaks are highlighted in blue (padj<0.05, fold change>1.5 or <-1.5). **D.** Motif analysis of the regions with increased CDK8 DNA binding peaks from CUT&RUN analysis in Rh30 cells. **E.** Heatmaps showing chromatin occupancy of CDK8, CCNC, MED12, and MED13 at regions with upregulated SIX4 binding at 24 hrs of DMSO or BI-1347 treatment. **F.** IGV gene tracks showing the PRO-seq, CDK8, CCNC, MED12, and MED13 binding at the *RUNX1* gene body and enhancer loci at indicated time points after BI-1347 treatment. **G.** Heatmaps of CDK8, CCNC, MED12, and MED13 CUT&RUN signal around PAX3::FOXO1-regulated enhancers before and after 24 hrs of BI-1347 treatment. **H.** IGV gene tracks showing the binding of CDK8, CCNC, MED12, and MED13 at a *RUNX2* super enhancer cluster at indicated time points after BI-1347 treatment.

Lastly, we investigated whether the chromatin binding of Mediator kinase module components was altered with CDK8 inhibitor treatment. We performed CUT&RUN with antibodies against CDK8, CCNC, MED12, or MED13 after BI-1347 treatment for 24 hours. As expected, there were no significant global changes in chromatin binding of CDK8, CCNC, MED12 or MED13 (**Fig. 7C, S6E-S6G**). However, we observed specific sites where binding of the CDK8 kinase module components was increased (**Fig. 7C, S6E, S6F**). Notably, Homer motif analysis revealed that CDK8-bound regions with increased occupancy upon CDK8 inhibition are enriched for motifs associated with myogenic differentiation such as MyoG and RUNX1 (**Fig. 7D**). Moreover, at SIX4 upregulated loci, we detected a substantial increase in chromatin occupancy of all four kinase module components, a pattern that was similarly observed at TADA2B upregulated sites (**Fig. 7E, S6H**). For instance, at the *RUNX1* locus, the upstream enhancer and gene body region showed active transcription, accompanied by increased Mediator kinase components to the enhancer (**Fig. 7F**). In contrast, no changes in Mediator kinase component binding were observed at PAX3::FOXO1-regulated core enhancers, such as a *RUNX2* super enhancer loci (**Fig. 7G, 7H**). These findings suggest that CDK8 inhibition does not broadly alter chromatin occupancy of the Mediator kinase module. Rather, CDK8 small molecule inhibitors selectively increase Mediator kinase, SAGA complexes, and SIX4 on chromatin at differentiation-associated enhancers, facilitating transcriptional activation of myogenic genes and contributing to the pro-differentiation effects of CDK8 inhibition in aRMS.

## DISCUSSION

Previous studies established that CDK8 functions as a transcriptional repressor by modulating RNA polymerase II (Pol II) activity or by phosphorylating transcription factors^26,30,40^. In yeast, Mediator complexes with and without the CDK8 kinase module were identified, with the majority of Pol II and MED26-containing complexes lacking the CDK8 module^41–43^. This modularity highlights CDK8 as a reversible molecular switch that fine-tunes transcription reinitiation by restricting Pol II interactions with the core Mediator complex^26,44,45^. Furthermore, CDK8 has been shown to inhibit the kinase activity of general transcription initiation factor IIH, thereby modulating transcription initiation^46^. This enhancer-Mediator-Pol II interplay is crucial for regulating transcriptional programs, as demonstrated in naïve pluripotent stem cells, where CDK8 inhibition activates pluripotency-associated transcription^47–49^. Similarly, this principle is also demonstrated in AML, where super-enhancer associated genes are up regulated upon CDK8 inhibition^48^.

Our findings provide compelling evidence that the CDK8 inhibitor BI-1347 releases transcriptional repression in aRMS, where CDK8 is more highly expressed compared to human skeletal muscle cells and other types of tumors, primarily by derepressing differentiation-related transcriptional programs. These observations align with previous studies showing that CDK8 represses differentiation pathways to sustain the oncogenic properties of stem- or progenitor-like cells in various cancers^50,51^. Phenotypically, we observed a proliferation defect and promotion of differentiation after CDK8 inhibition, findings supported by the upregulation of genes associated with myogenic differentiation, such as *VGLL2*, *RUNX1,* and *SEMA3D*. This aligns with findings in AML, where CDK8 inhibition promoted cellular maturation, and in colon cancer, where it induced differentiation^14,48^. In the case of aRMS, both CDK8 inhibition and PAX3::FOXO1 degradation release the block on muscle differentiation imposed by PAX3::FOXO1, allowing aRMS cells to adopt a more differentiated phenotype. The transcriptional reprogramming influences the expression of genes involved in myogenic differentiation pathways and underpins the shift of aRMS cells from a proliferative state to a more differentiated state after CDK8 inhibition.

Previous studies report that PAX3::FOXO1 recruits transcriptional regulatory factors, including CDK8 and SMARCA4, and that CDK8 localization on chromatin is decreased with the reduced chromatin accessibility following rapid PAX3::FOXO1 degradation^14,52^. Whether these transcriptional regulatory factors are required for the chromatin binding of PAX3::FOXO1 broadly or its target sites is not well defined. Our data suggests that CDK8 inhibition does not broadly impact PAX3::FOXO1 chromatin occupancy or its transcriptional regulatory function. Even at the PAX3::FOXO1 controlled core enhancers, chromatin association of PAX3::FOXO1 and CDK8 remained intact. Notably, CDK8 inhibition selectively activated differentiation-related enhancers without affecting “stemness” transcription factors directly regulated by PAX3::FOXO1 at early time points. The transcriptional profiles induced by CDK8 inhibition and PAX3::FOXO1 degradation, however, showed partial overlap, particularly in the activation of myogenic differentiation genes at later time points, suggesting that the secondary effects of these perturbations are related. And, while the examination of PAX3::FOXO1 protein levels and its sub-cellular localization does not support a direct regulation of PAX3::FOXO1 by CDK8, we cannot rule out the possibility of phosphorylation of the fusion or other transcription factors by CDK8. In addition, the study of SMARCA4 suggests that the degradation of SMARCA4 also induces myogenic differentiation in aRMS^52^. Whether Mediator and BAF complex cooperate in PAX3::FOXO1 aRMS to regulate transcription remains to be determined.

Our genome-scale CRISPR BI-1347 anchor screen identified the critical role of SAGA complex in mediating the transcriptional coactivation and differentiation-promoting effects of CDK8 inhibition in aRMS. Specifically, the identification of the knockout of SAGA components TADA2B and KAT2A from the histone acetyltransferase (HAT) module and TAF5L from the core module as key resistance hits in the genome-scale CRISPR-Cas9 screen highlights the essential collaboration between Mediator and SAGA in the regulation of differentiation-associated transcription. Disruption of these proteins attenuated both the transcriptional activation of differentiation-related enhancers at genes, such as *RUNX1, VGLL2,* and *SEMA3D,* and growth inhibition induced by CDK8 inhibitors, suggesting that SAGA’s HAT activity facilitates the enhancer-driven transcriptional elongation required for muscle differentiation. While at baseline, TADA2B is localized at both promoters and enhancers, the increased TADA2B with CDK8 inhibition was at sites with increased open chromatin at enhancers. These findings align with our observation of increased enhancer RNA transcription and chromatin accessibility at differentiation-associated loci following CDK8 inhibition.

The CRISPR screen also identified a muscle development related transcription factor needed for the full activity of BI-1347. SIX4 and other SIX family member proteins play an essential role in several stages of skeletal muscle development. While SIX1 is reported to repress RMS differentiation, SIX4 is reported to promote the early specification of hypaxial myogenic progenitors through transactivation of PAX3, while later controlling myoblast determination and differentiation via activation of *MYF4, MYOD1, MYF6*, and *MYOG*^38,53–55^. In aRMS, where PAX3::FOXO1 enforces an undifferentiated state, our findings suggest that the CDK8 inhibitor BI-1347 disrupts this oncogenic program in part by inducing a differentiation permissive transcriptional landscape, leading to the activation of SIX4-driven muscle differentiation programs. In our screen, other upregulated muscle differentiation-related genes, such as *SEMA3D, RUNX2,* and *VGLL2,* do not score as rendering resistance to BI-1347 when knocked out, suggesting they may act as downstream targets of SIX4-induced myogenic differentiation transcriptional cascade rather than the upstream node.

While there is a strong correlation between aRMS dependency on *CDK8* by CRISPR knockout and the response to CDK8 chemical inhibition, we noted that pharmacological inhibition of CDK8 resulted in more dynamic transcriptional changes compared to genetic knockout. Moreover, in addition to identifying members of the SAGA complex and SIX4, the BI-1347 CRISPR modifier screen also identified that knockout of Mediator kinase members rendered partial resistance to BI-1347. Finally, we determined that there is a gain of binding at specific sites on chromatin of Mediator kinase module members, such as CDK8, CCNC, MED12, and MED13, particularly at the sites where SIX4 and SAGA complex are also gained with CDK8 inhibitor treatment in aRMS cells. All told, these results support the hypothesis that BI-1347 may have both a loss of function (inhibition of CDK8 enzymatic activity) and gain of function effect (inhibitor trapping) on Mediator regulated transcription, which will be the focus of our future studies. Kinase inhibitor trapping has been observed in other studies. Structural analyses showed that imatinib becomes stably entrapped within the ABL kinase, and a similar kinase trapping mechanism has been reported for p38a kinase inhibitor, resulting in enhanced compound affinity^56^. Similarly, PARP inhibitors work in part by trapping PARP at sites of DNA damage preventing its repair^57^. Here, our data suggests that CDK8 inhibition and the trapping of inhibited mediator kinase enhances the activity of the mediator complex.

Induction of myogenic differentiation in aRMS by the CDK8 inhibitor BI-1347 suggests that CDK8 inhibitors may serve as modulators to counteract aRMS oncogenic activity. By disrupting the repressive transcriptional network maintained by PAX3::FOXO1 and inducing the activated transcriptional network maintained by other transcription factors such as SIX4, CDK8 inhibitors can restore differentiation programs, offering a novel therapeutic strategy for aRMS. Indeed, the use of differentiation therapy has expanded in recent years with the FDA approval of IDH and menin inhibitors for the treatment of AML^58,59^. In considering the translation of CDK8 inhibition to the clinic, early safety concerns were raised as the CDK8-/-mouse is embryonic lethal with failure of compaction of embryonic blastomeres ^60^. CDK8, however, is not essential for cell growth or survival of *Drosophila* S2 cells or 293T cells, and systemic inducible *CDK8* knockout mice have no survival defect^60–62^. Encouragingly, CDK8 inhibitors such as SEL-120-34A have shown promise in preclinical studies and are currently in clinical trials for treatment of advanced AML and high-risk myelodysplastic syndrome^31^. In summary, our findings highlight the potential of CDK8 inhibitors as differentiation-inducing agents in aRMS and reveal new insights about the mechanistic role of CDK8 in this disease.

## METHODS

### Data Availability

This paper does not report original software for NGS data analysis. Any code for analysis or data visualization will be shared upon request.

### Cell Lines

The Rh28 cell line was cultured in RPMI-1640 (Corning by Mediatech, Inc.) containing 10% FBS (Sigma Aldrich). Rh30_PAX3::FOXO1-FKBP-HA cell lines weas cultured in RPMI-1640 (Corning by Mediatech, Inc.) containing 10% FBS (Sigma Aldrich), and was generously provided by Dr. Scott Hiebert at Vanderbilt University, Nashville, TN. Rh30, Rh4, RHJT, and HEK293T cell lines were cultured in DMEM (Corning by Mediatech, Inc.) containing 10% FBS (Sigma Aldrich). SkMC cell line was cultured in skeletal muscle cell growth media (PromoCell) with 5% FBS (Sigma Aldrich). All cell lines were consistently tested mycoplasma negative (MycoAlert, Lonza). All cell lines were confirmed with correct identities by short tandem repeats (STR) profiling.

### Small Molecular Inhibitors

BI-1347 and BI-1374 were generously provided by Boehringer Ingelheim through OpnMe.com. SEL-120-34A was purchased from Sellechchem (#S8840). JH-XII-178 was generously provided by Dr. John Hatcher and Dr. Nathanael Gray. GSK699 was synthesized by WuXi AppTec.

### Small harpin RNAs (shRNA)

The shRNA was cloned into pLOK.1-puro (Addgene, plasmid#8453) and Tet-pLKO-puro (Addgene, plasmid#21915).

shNT: CCGGCAACAAGATGAAGAGCACCAACTCGAG TTGGTGCTCTTCATCTTGTTG TTTTTG

shCDK8-1: CCGGATGTCCAGTAGCCAAGTTCCACTCGAGTGGAACTTGGCTACTGGACATTTTT T

shCDK8-2: CCGGCGAGAGCTTAAGCATCCAAACCTCGAGGTTTGGATGCTTAAGCTCTCGTTTT TTG

### Single-guide RNAs (sgRNA)

The sgRNA was cloned into lentiCRISPR v2 containing Cas9 and puromycin resistance marker plasmid (Addgene, plasmid#52961).

sgChr2.2: GGTGTGCGTATGAAGCAGTG

sgCDK8-3: GAGGACCTGTTTGAATACGA

sgCDK8-4: AGTGACTTCACCATTCCCCG

sgTADA2B-1: GACAGGTGTGGTCTGTCACG

sgTADA2B-3: GGGCGGCTGGACCAGTCGCG

sgTAF5L-1: AAACAGTCCGAAGAGCACAG

sgTAF5L-3: GCAGAAACATTCCATGGAAG

sgSIX4-1: GCCTGCCCAGAAGTTCCGAG

sgSIX4-2: GGATCAGGTACAACTCCACT

### Western Blot

Cells were collected at indicated timepoints and lysed with RIPA buffer (50nM Tris pH8.0, 150mM NaCl, 1% NP-40, 0.5% sodium deoxycholate, 0.1% SDS) containing protease inhibitors (Aprotinin, Sigma Aldrich #6279; Phenylmethanesulfonyl fluoride solution, Sigma Aldrich #93482; Phosphatase Inhibitor Cocktail 3, Sigma Aldrich #P0044). Lysates were sonicated and cleared by centrifugation, and then subjected to NuPAGE Tris-Acetate protein gels to separate proteins (ThermoFisher, #EA03785BOX) and electrophoretic transfer to membranes. Primary antibodies used in this study includes: CDK8 (ProteinTech, #22067; Cell Signaling Technology, #4106S; Santa Cruz Biotechnology, #SC-13155; abcam, #ab229192), STAT1 (Cell Signaling Technology, #9176S), pSTAT1S727(Cell Signaling Technology, #8826S), SIX4 (Santa Cruz Biotechnology, #SC-390779), GAPDH (Santa Cruz Biotechnology, #SC-365062), LaminB (Abcam, #ab16048; Santa Cruz Biotechnology, #SC-374015), ACTB (Sigma Aldrich, #A2066), full PARP (Cell Signaling Technology, #9542S), cleaved PARP (Cell Signaling Technology, #5625S), PAX3::FOXO1 (CancerTools, #160866), MYC (XXX), streptavidin (XXX), TAF5L (ProteinTech, #19274), TADA2B (ProteinTech, #17367), KAT2A (ProteinTech, #66575), KAT2B (Cell Signaling Technology, #C14G9), CCNC (ProteinTech, #26464), MED13 (ProteinTech, #26464), MED12 (ProteinTech, #20028). Signal was visualized with secondary HRP-linked antibodies (Cell Signaling Technology, #7074S; Cell Signaling Technology, #7076S) using the Licor Odyssey imaging system, or IR-Dye conjugated antibodies (Fisher Scientific, #NC0250902; Fisher Scientific, #NC0250903;) using ECL western blot substrates (Fisher Scientific, #54002401) on X-ray film (Life Technologies, #34089).

### Cell Growth Assays

Trypan Blue exclusion assay: Trypan Blue exclusion assay: Cells were transduced with shNT, shCDK8-1, or shCDK8-2, and seeded at 2×10^5^ cells/well after puromycin selection (3 days of antibiotic selection). Cells were then stained with Trypan Blue Solution (Sigma Aldrich, #T8154) followed by automated cell counting on a hematocytometer (Bio-Rad TC20 automated cell counter). Viable cells were counted with Trypan Blue dye exclusion every day for 4 days consecutive. Quantification was performed in triplicate and the values averaged and shown with standard deviations. On-target editing of the shRNA was confirmed by Western blot.

CellTiter-Glo (CTG) cell viability assay: Cells were treated with indicated small molecule inhibitors or were transfected with indicated sgRNA were seeded into 384-well plates at 800 cells/well. CellTiter-Glo Luminescent assay (Promega, #G7573) were performed according to the manufacturer’s instructions. Each condition includes at least 6 replicates, and each experiment were at least triplicate. On-target editing of the CRISPR guides and small molecule inhibitors was confirmed by Western blot.

IncuCyte Live cell proliferation assay: Cells with indicated sgRNA transfection, small molecule inhibitors treatment, or combination were seeded in 96-well plates at 2,000 cells/well and allowed to adhere overnight. Images were captured every 12 hours over the course of the experiment using phase-contrast imaging. Confluence was quantified using the IncuCyte integrated analysis software, and proliferation curves were generated based on percent confluence over time. All conditions were assessed in technical triplicates and normalized to control treatments as appropriate.

### *In vivo* Xenografts Tumor Experiment

Rh28 cells with CDK8-targeting shRNAs or Rh30 cells were harvested, counted, and resuspended in a 1:1 mixture of PBS and Matrigel (Corning, #354234). A total of 1×10^7^ cells were subcutaneously injected into the flanks of Fox Chase SCID Beige mice (Charles River Laboratories, #250). To induce shRNA expression, mice were fed with doxycycline-containing chow (Envigo, #TD.01306) starting on the day of established tumors (∼100 mm^3^). For CDK8 inhibitor SEL-120-34A study, mice were treated with SEL-120-34A by oral gavage at 40 mg/kg daily. Animal weight was monitored, and tumor size was measured 3 times a week using digital calipers. Mice were euthanized at endpoint, and tumors were collected for downstream analysis. All animal procedures were performed in accordance with institutional IACUC guidelines.

### Cell Cycle Analysis

Cells were treated with DMSO or BI-1347 for 3, 7, 14, 21 days, and pulsed with 20μM - bromo-2’-deoxyuridine (BrdU) for 2 hours and fixed overnight with 70% ethanol at 4°C. After staining with fluorescein isothiocyanate (FITC)-conjugated anti-BrdU (BD Biosciences, #556028) and counterstaining with 7-AAD (BD Biosciences, #559925), cells were analyzed by flow cytometry. All flow cytometry figures were generated using FlowJo software (v10.9).

### Apoptosis Analysis

Annexin V Assay: Cells were treated with DMSO or BI-1347 for 3, 7, 14, 21 days and 1×10^5^ cells were collected for FITC-conjugated Annexin V and propidium iodide (PI) staining (BD Biosciences, #556547). Cells were analyzed by flow cytometry. All flow cytometry figures were generated using FlowJo software (v10.9).

Caspase-Glo® 3/7 Assay: Cells were transduced with shNT, shCDK8-1, or shCDK8-2, and cells were seeded at 2×10^4^ cells/well in 96-well plate after 3 days of antibiotic selection. After 4 days, 10ul of Caspase-Glo reagent was added to each well, and plates were incubated at room temperature for 30 minutes before measuring luminescent signal. Each condition includes at least 6 replicates, and each experiment were at least triplicate.

### Immunofluorescence

Cells were seeded on coverslips and treated with DMSO or BI-1347 for 7 days. After fixation using 3.7% paraformaldehyde for 15 minutes at room temperature, the cells were permeabilized using 0.3% Triton X-100 for 30 minutes. Cells were blocked with 1% BSA in PBS for 30 minutes in a humidified chamber, and then incubated with primary Myogenin antibody (Abcam, #ab1835) diluted in blocking buffer containing 0.5% NP-40 for 1 hour at 37°C. After 3 times PBS washing, cells were incubated in Alexa Fluor secondary antibody (Abcam ab150117, Invitrogen A-11034) with Phalloidin (Thermo Fisher Scientific, A12380) and DAPI diluted in blocking buffer containing 0.5% NP-40 in PBS at 37 °C for 45 minutes. Coverslips were mounted on slides with Prolong Gold Antifade reagent (Thermo Fisher Scientific, P36930) and dried overnight in the dark. Images were collected using a Nikon fluorescent microscope.

### Immunohistochemistry

A rhabdomyosarcoma patient-derived xenograft (PDX) microarray was obtained from St. Jude Children’s Research Hospital, courtesy of Dr. Elizabeth Stewart. The microarray sample sections were deparaffinized, rehydrated, and subjected to antigen retrieval in antigen unmasking buffer (Vector Labs, #H-3300). Endogenous peroxidase activity was blocked with 3% hydrogen peroxide, followed by 30 minutes blocking buffer incubation at room temperature. Sections were incubated overnight at 4°C with an anti-CDK8 primary antibody (Santa Cruz, #sc-13155), followed by a biotinylated secondary antibody (Vector Labs #PK-6200) and detection using a DAB chromogen kit (Vector Labs #SK-4100). Slides were counterstained with hematoxylin (Sigma Aldrich, #MHS16), dehydrated, and mounted.

### Hematoxylin and Eosin (H&E) Staining

Sections of FFPE xenograft tumors tissue were deparaffinized in xylene, rehydrated through a graded series of ethanol, and rinsed in distilled water. Slides were stained with hematoxylin to visualize nuclei, followed by differentiation in acid alcohol and bluing in ammonium hydroxide. Eosin was then applied to counterstain cytoplasmic and extracellular components. After dehydration and clearing, sections were mounted with coverslips and visualized under a light microscope. Tumor samples were submitted to the Duke Pathology Core for analysis.

### TaqMan Real-Time PCR Assay

Cells with indicated treatment conditions were collected and total RNA was extracted using TRIzol reagent (Life Technologies, #15596026) and reverse-transcribed into cDNA using the High-Capacity cDNA Reverse Transcription Kit (Life Technologies, #1708891). Real-time PCR reactions were carried out in 384-well plates using TapMan Universal PCR Master Mix (Life Technologies, #4374338) on a QuantStudio 6 Flex Real-Time PCR System under standard cycling conditions. Gene expression was quantified using the ΔΔCt method, with GAPDH as an internal control. All reaction were run in six technical replicates, and relative expression was normalized to control samples. TapMan probes in this study:

GAPDH: Life Technologies, #Hs02786624_g1

RUNX1: Life Technologies, #Hs02558380_m1

SEMA3D: Life Technologies, #Hs00380877_m1

VGLL2: Life Technologies, #Hs00403461_m1

### Precision Nuclear Run-on and Sequencing (PRO-seq) and Data Processing

PRO-seq was performed in biological replicates as previously described using 10 million cells treated with DMSO or BI-1347 for 4, 24, and 72 hours^63^. After washing and permeabilization, nuclei were stored in freezing buffer and then subject to biotin run-on using biotin-11-ATP, biotin-11-GTP, biotin-11-CTP, and biotin-11-UTP (Perkin Elmer, #NEL54[2,3,4,5]001). RNA was reversed transcribed and amplified to make the cDNA library for sequencing by the Harvard Medical School Nascent Transcriptomics Core. The sequences were aligned to hg38 using bowtie2 before using the Nascent RNA Sequencing Analysis (NRSA) pipeline^34^ to determine the gene body and eRNA changes.

### RNA Sequencing and Data Processing

All RNA-seq experiments were performed in biology triplicate. Cells with indicated treatment were collected and total RNA was extracted using TRIzol reagent. RNA was submitted to the Novogene for library preparation and sequencing. Gene expression values were derived from paired end RNA-Seq data. The RNA-seq processing pipeline was roughly modeled on the GTEx pipeline (https://github.com/broadinstitute/gtex-pipeline/). FastQC was used to evaluate read quality on raw RNA-seq reads. Reads were aligned to the human genome (hg38) using STAR (v2.7.2b) (Dobin et al., 2013). Transcript-level quantifications were calculated using RSEM (v1.3.1) (Li et al. 2011). Gene counts from STAR were then used to quantify differentially expressed genes between the experimental and control conditions using the R Bioconductor package DESeq2 (Anders, et al., 2010) using the approximate posterior estimation for GLM coefficients (apeglm) method for effect size. Normalized expression values for individual samples were obtained from RSEM log2 TPM values with the RSEM log2(TPM+1) values used for GSEA and producing RNA-seq heat map plots.

### Cleavage Under Targets and Release Using Nuclease (CUT&RUN) and Data Processing

CUT&RUN experiments were performed in biological duplicate as described^64^. In Brief, 250,000 cells with indicated treatment were collected and incubated with CUTANA Concanavalin A conjugated paramagnetic beads (EpiCypher, #21-1401) for 10 minutes at room temperature. Cells were permeabilized using 0.02% freshly dissolved digitonin (Sigma Aldrich, #300410), and then subject to primary antibodies incubation at 4°C overnight. Primary antibodies used for CUT7RUN in this study includes: CDK8 (ProteinTech, #22067), SIX4 (Santa Cruz Biotechnology, #SC-390779), HA (Cell Signaling Technology, #C29F4-3724), TADA2B (ProteinTech, #17367), CCNC (ProteinTech, #26464), MED13 (ProteinTech, #26464), MED12 (ProteinTech, #20028), H3K4me3 (EpiCypher, #13-0041), H3K27ac (Cell Signaling Technology, #8173S). Following washing, targeted regions of the genome were digested using CUTANA pAG-MNase (EpiCypher, #15-1116) fusion protein with 2 hours incubation at 0 °C. DNA was then extracted using Monarch PCR and DNA Cleanup Kit (New England Biolabs, #T1030S), and the sequencing libraries were generated using the NEBNext Ultra II DNA Library Prep Kit (New England Biolabs, #E7654L) and NEBNext Multiplex Oligos for Illumina 96 Unique Dual Index Primer Pairs (New England Biolabs, #E6440S). Sequencing was performed by the Molecular Biology Core Facilities (MBCF) at Dana-Farber Cancer Institute (DFCI).

Sequencing read quality was examined using FastQC (http://www.bioinformatics.babraham.ac.uk) (v0.11.9). Raw FASTQ data were trimmed using Trimmomatic-v0.38 and paired end reads were aligned to human hg38 genome using Bowtie2 (v2.3.5) in very sensitive local mode (--phred33 -I 0 -X 2000 -N 1 --very-sensitive). Peaks were called using MAXS2 (v2.1.4) with a threshold of q<0.01 and peak were annotated to the nearest TSS using HOMER. Differential analysis was performed using DiffBind and DESeq2. Significantly changed peaks were defined by a 1.5-fold change threshold and FDR<0.05. Bigwig files were normalized and generated using Deeptools. Heatmaps were created by Deeptools using the normalized bigwig files.

### Assay for Transposase Accessible Chromatin using Sequencing (ATAC-seq) and Data Processing

ATAC-seq was performed in biological triplicate following the manufacturer’s protocol (Active Motif, #53150). 100,000 cells with indicated treatment were used to isolate nuclei. Nuclei were incubated with tagmentation Master Mix at 37 °C for 30 minutes after lysing the cells in ice-cold ATAC Lysis Buffer and the DNA was purified with the DNA purification column. PCR amplification of tagmented DNA was performed to make libraries with the appropriate indexed primers. After SPRI bead clean-up, the DNA libraries were submitted to the Molecular Biology Core Facilities (MBCF) at Dana-Farber Cancer Institute (DFCI) for sequencing.

Raw FASTQ data were trimmed by Trimmomatic-v0.39, and paired end reads were aligned to a concatenated human genome hg38 using Bowtie2 (v2.3.5) (--phred33 -I 0 - X 2000 -N 1 --very-sensitive). Peaks were called using Genrich (v. 0.6.1) (https://github.com/jsh58/Genrich) with the following options -j -y -r -e chrM -q 0.05 -a 20. Peaks were annotated to the nearest TSS using HOMER. Differential analysis was performed using DiffBind and DESeq2. Significantly changed peaks were defined by a 1.5-fold change threshold and FDR<0.05. Bigwig files were generated and normalized using the DESeq2 size factors using Deeptools. Heatmaps were created by Deeptools using the normalized bigwig files.

### Bio-ID Labeling and Mass Spectrometry

For each replicate, approximately 7×10^6^ cells were transfected using Fugene 6 (Promega) transfection reagent as according to manufacturer’s protocol and grown in DMEM. 48 hours later biotin was added to the cultures at 50 µM and cultured for an additional 16 hours. Each replicate was lysed in 200 µL RIPA supplemented with Complete protease inhibitors (Roche), rotated end-over-end for 30 min and cleared by centrifugation at 4° C. MydOne streptavidin coated agarose beads (Invitrogen) were simultaneously equilibrated in RIPA buffer. 90% of lysate was then added to the beads and rotated end-over-end for 16 hrs at 4° C. The remaining 10% was used as input. Beads were then washed 5 times in wash buffer (0.5% (w/v) sodium deoxycholate, 0.5% (w/v) igepal CA-630, 1 mM EDTA, 100 mM NaCl, 50 mM Tris.Cl, pH 7.5) by end-over-end incubation for 5 minutes at 4° C. After final wash, beads were resuspended in 40ul PBS + 2.5 uM Biotin, and 10 ul 5× SDS Page loading buffer. 40 µL of each sample was loaded onto an Invitrogen NuPAGE 4-12% SDS-PAGE gel and run for approximately 5 min to electrophorese all proteins into the gel matrix. The entire MW range was then excised in a single gel-band and subjected to standardized in-gel reduction, alkylation, and tryptic digestion. Following lyophilization of the extracted peptide mixtures, samples were resuspended in 20 µL of 2% acetonitrile/1% TFA supplemented with 12.5 fmol/µL yeast ADH. From each sample, 3 µL was removed to create a QC Pool sample which was run periodically throughout the acquisition period.

Quantitative LC/MS/MS was performed on 4 µL of each sample, using a nanoAcquity UPLC system (Waters Corp) coupled to a Thermo QExactive HF-X high resolution accurate mass tandem mass spectrometer (Thermo) via a nanoelectrospray ionization source. Briefly, the sample was first trapped on a Symmetry C18 20 mm × 180 µm trapping column (5 μl/min at 99.9/0.1 v/v water/acetonitrile), after which the analytical separation was performed using a 1.8 µm Acquity HSS T3 C18 75 µm × 250 mm column (Waters Corp.) with a 90-min linear gradient of 5 to 30% acetonitrile with 0.1% formic acid at a flow rate of 400 nanoliters/minute (nL/min) with a column temperature of 55C. Data collection on the QExactive HF mass spectrometer was performed in a data-dependent acquisition (DDA) mode of acquisition with a r=120,000 (@ m/z 200) full MS scan from m/z 375 – 1600 with a target AGC value of 3e6 ions followed by 30 MS/MS scans at r=15,000 (@ m/z 200) at a target AGC value of 5e4 ions and 45 ms. A 20s dynamic exclusion was employed to increase depth of coverage. The total analysis cycle time for each sample injection was approximately 2 hours. Following UPLC-MS/MS analyses, data was imported into Rosetta Elucidator v 4.0 (Rosetta Biosoftware, Inc.), and analyses were aligned based on the accurate mass and retention time of detected ions (“features”) using PeakTeller algorithm in Elucidator. Relative peptide abundance was calculated based on area under-the-curve (AUC) of the selected ion chromatograms of the aligned features across all runs. The MS/MS data was searched against the SwissProt H. sapiens database (downloaded in Aug 2017) with additional proteins, including yeast ADH1, bovine serum albumin, as well as an equal number of reversed-sequence “decoys”) false discovery rate determination. Mascot Distiller and Mascot Server were utilized to produce fragment ion spectra and to perform the database searches. Database search parameters included fixed modification on Cys (carbamidomethyl) and variable modifications on Meth (oxidation) and Asn and Gln (deamidation). After individual peptide scoring using the PeptideProphet algorithm in Elucidator, the data was annotated at a 0.8% peptide false discovery rate.

### CRISPR-Cas9 Drug Modifier Screen and Data Processing

A genome-wide CRISPR-Cas9 loss-of-function screen was performed in Rh30 cells treated with a sublethal concentration (∼IC50) of CDK8 inhibitor BI-1347 to identify genetic modifiers of sensitivity or resistance to BI-1347 response. Cells stably expressing Cas9-mCherry were transduced with the genome-wide sgRNA library Avana-4 in duplicate at a low multiplicity of infection (MOI∼0.3) to ensure single sgRNA integration per cell, followed by puromycin selection. After recovery, cells were split into two arms and treated with either DMSO or BI-1347 at a concentration sufficient to induce partial growth inhibition (IC30) for 21 days. Cells were collected at day-14 and day-21, and genome DNA was extracted (Takara Bio # 740950.50). Genomic DNA was submitted to the Broad Institute Genetic Perturbation Platform (GPP) for PCR-amplification and sequencing. sgRNA representation and gene-level enrichment were analyzed using usingthe Broad Institute of MIT and Harvard Genetic Perturbation Platform’s (GPP) APRON analysis tool (https://portals.broadinstitute.org/gppx/apron/screener/) to identify candidate genes whose loss sensitized or conferred resistance to BI-1347 treatment.

## Acknowledgements

This work was funded by support from National Cancer Institute R35 CA283977 (K.S.), U54 CA231630 (C.M.C, C.M.L., K.C.W.); the CureSearch for Children’s Cancer Foundation, the Rally Foundation for Childhood Cancer Research, and the St. Baldrick’s Foundation (C.M.L.); the Palumbo Rhabdomyosarcoma Research Fund, and the Corinne Cecilia Sciarappa Fund. S.Z. is supported by an Alex’s Lemonade Stand Fellowship (#22-27271). M.J. and B.G. were supported by NCI T32HD094671 Fellowships.

This work was supported by the Pediatric Cancer Dependencies Accelerator of the Broad Institute, Dana-Farber Cancer Institute, and St. Jude Children’s Research Hospital: pdedep.org

## REFERENCES

1. Mitelman F, Johansson B, Mertens F. The impact of translocations and gene fusions on cancer causation. Nat Rev Cancer. Apr 2007;7(4):233–45. doi:10.1038/nrc2091

2. Lobato MN, Metzler M, Drynan L, Forster A, Pannell R, Rabbitts TH. Modeling chromosomal translocations using conditional alleles to recapitulate initiating events in human leukemias. J Natl Cancer Inst Monogr. 2008;(39):58–63. doi:10.1093/jncimonographs/lgn022

3. Mertens F, Antonescu CR, Mitelman F. Gene fusions in soft tissue tumors: Recurrent and overlapping pathogenetic themes. Genes Chromosomes Cancer. Apr 2016;55(4):291–310. doi:10.1002/gcc.22335

4. Sorensen PH, Lynch JC, Qualman SJ, et al. PAX3-FKHR and PAX7-FKHR gene fusions are prognostic indicators in alveolar rhabdomyosarcoma: a report from the children’s oncology group. J Clin Oncol. Jun 01 2002;20(11):2672–9. doi:10.1200/JCO.2002.03.137

5. Ognjanovic S, Linabery AM, Charbonneau B, Ross JA. Trends in childhood rhabdomyosarcoma incidence and survival in the United States, 1975-2005. Cancer. Sep 15 2009;115(18):4218–26. doi:10.1002/cncr.24465

6. Ma X, Liu Y, Alexandrov LB, et al. Pan-cancer genome and transcriptome analyses of 1,699 paediatric leukaemias and solid tumours. Nature. Mar 15 2018;555(7696):371–376. doi:10.1038/nature25795

7. Que Y, Wang J, Sun F, et al. Safety and clinical efficacy of sintilimab (anti-PD-1) in pediatric patients with advanced or recurrent malignancies in a phase I study. Signal Transduct Target Ther. Oct 13 2023;8(1):392. doi:10.1038/s41392-023-01636-9

8. Ciurej A, Lewis E, Gupte A, Al-Antary E. Checkpoint Immunotherapy in Pediatric Oncology: Will We Say Checkmate Soon? Vaccines (Basel). Dec 12 2023;11(12)doi:10.3390/vaccines11121843

9. Davis KL, Fox E, Merchant MS, et al. Nivolumab in children and young adults with relapsed or refractory solid tumours or lymphoma (ADVL1412): a multicentre, open-label, single-arm, phase 1-2 trial. Lancet Oncol. Apr 2020;21(4):541–550. doi:10.1016/S1470-2045(20)30023-1

10. Geoerger B, Kang HJ, Yalon-Oren M, et al. Pembrolizumab in paediatric patients with advanced melanoma or a PD-L1-positive, advanced, relapsed, or refractory solid tumour or lymphoma (KEYNOTE-051): interim analysis of an open-label, single-arm, phase 1-2 trial. Lancet Oncol. Jan 2020;21(1):121–133. doi:10.1016/S1470-2045(19)30671-0

11. Geoerger B, Zwaan CM, Marshall LV, et al. Atezolizumab for children and young adults with previously treated solid tumours, non-Hodgkin lymphoma, and Hodgkin lymphoma (iMATRIX): a multicentre phase 1-2 study. Lancet Oncol. Jan 2020;21(1):134–144. doi:10.1016/S1470-2045(19)30693-X

12. Loeb DM, Lee JW, Morgenstern DA, et al. Avelumab in paediatric patients with refractory or relapsed solid tumours: dose-escalation results from an open-label, single-arm, phase 1/2 trial. Cancer Immunol Immunother. Oct 2022;71(10):2485–2495. doi:10.1007/s00262-022-03159-8

13. Kailayangiri S, Altvater B, Meltzer J, et al. The ganglioside antigen G(D2) is surface-expressed in Ewing sarcoma and allows for MHC-independent immune targeting. Br J Cancer. Mar 13 2012;106(6):1123–33. doi:10.1038/bjc.2012.57

14. Zhang S, Wang J, Liu Q, et al. PAX3-FOXO1 coordinates enhancer architecture, eRNA transcription, and RNA polymerase pause release at select gene targets. Mol Cell. Nov 11 2022;doi:10.1016/j.molcel.2022.10.025

15. Liu B, Liu X, Han L, et al. BRD4-directed super-enhancer organization of transcription repression programs links to chemotherapeutic efficacy in breast cancer. Proc Natl Acad Sci U S A. Feb 08 2022;119(6)doi:10.1073/pnas.2109133119

16. Gryder BE, Yohe ME, Chou HC, et al. PAX3-FOXO1 Establishes Myogenic Super Enhancers and Confers BET Bromodomain Vulnerability. Cancer Discov. Aug 2017;7(8):884–899. doi:10.1158/2159-8290.CD-16-1297

17. Lamonica JM, Deng W, Kadauke S, et al. Bromodomain protein Brd3 associates with acetylated GATA1 to promote its chromatin occupancy at erythroid target genes. Proc Natl Acad Sci U S A. May 31 2011;108(22):E159–68. doi:10.1073/pnas.1102140108

18. Tontsch-Grunt U, Traexler PE, Baum A, et al. Therapeutic impact of BET inhibitor BI 894999 treatment: backtranslation from the clinic. Br J Cancer. Aug 2022;127(3):577–586. doi:10.1038/s41416-022-01815-5

19. Krivtsov AV, Evans K, Gadrey JY, et al. A Menin-MLL Inhibitor Induces Specific Chromatin Changes and Eradicates Disease in Models of MLL-Rearranged Leukemia. Cancer Cell. Dec 09 2019;36(6):660–673.e11. doi:10.1016/j.ccell.2019.11.001

20. Kwon MC, Thuring JW, Querolle O, et al. Preclinical efficacy of the potent, selective menin-KMT2A inhibitor JNJ-75276617 (bleximenib) in KMT2A- and NPM1-altered leukemias. Blood. Sep 12 2024;144(11):1206–1220. doi:10.1182/blood.2023022480

21. Pan J, Meyers RM, Michel BC, et al. Interrogation of Mammalian Protein Complex Structure, Function, and Membership Using Genome-Scale Fitness Screens. Cell Syst. May 23 2018;6(5):555–568.e7. doi:10.1016/j.cels.2018.04.011

22. Malone CF, Mabe NW, Forman AB, et al. The KAT module of the SAGA complex maintains the oncogenic gene expression program in. Sci Adv. May 31 2024;10(22):eadm9449. doi:10.1126/sciadv.adm9449

23. Rzymski T, Mikula M, Żyłkiewicz E, et al. SEL120-34A is a novel CDK8 inhibitor active in AML cells with high levels of serine phosphorylation of STAT1 and STAT5 transactivation domains. Oncotarget. May 16 2017;8(20):33779–33795. doi:10.18632/oncotarget.16810

24. Hatcher JM, Vatsan PS, Wang E, Jiang J, Gray NS. Development of Highly Potent and Selective Pyrazolopyridine Inhibitor of CDK8/19. ACS Med Chem Lett. Nov 11 2021;12(11):1689–1693. doi:10.1021/acsmedchemlett.1c00300

25. DepMap, Broad (2024). DepMap 24Q2 Public. Figshare+. Dataset. 10.25452/figshare.plus.25880521.v1.

26. Tsai KL, Sato S, Tomomori-Sato C, Conaway RC, Conaway JW, Asturias FJ. A conserved Mediator-CDK8 kinase module association regulates Mediator-RNA polymerase II interaction. Nat Struct Mol Biol. May 2013;20(5):611–9. doi:10.1038/nsmb.2549

27. Zhang H, Chen DH, Mattoo RUH, et al. Mediator structure and conformation change. Mol Cell. Apr 15 2021;81(8):1781-1788.e4. doi:10.1016/j.molcel.2021.01.022

28. Tsai KL, Yu X, Gopalan S, et al. Mediator structure and rearrangements required for holoenzyme formation. Nature. Apr 13 2017;544(7649):196–201. doi:10.1038/nature21393

29. Treehouse Childhood Cancer Initiative. Treehouse Tumor Compendium V11 Public PolyA Dataset. UCSC Genomics Institute; 2023. Available from: https://treehousegenomics.soe.ucsc.edu/public-data/. 2023.

30. Bancerek J, Poss ZC, Steinparzer I, et al. CDK8 kinase phosphorylates transcription factor STAT1 to selectively regulate the interferon response. Immunity. Feb 21 2013;38(2):250–62. doi:10.1016/j.immuni.2012.10.017

31. U.S. N. National Library of Medicine. https://clinicaltrials.gov.

32. Kwak H, Fuda NJ, Core LJ, Lis JT. Precise maps of RNA polymerase reveal how promoters direct initiation and pausing. Science. Feb 22 2013;339(6122):950–3. doi:10.1126/science.1229386

33. Mahat DB, Kwak H, Booth GT, et al. Base-pair-resolution genome-wide mapping of active RNA polymerases using precision nuclear run-on (PRO-seq). Nat Protoc. Aug 2016;11(8):1455–76. doi:10.1038/nprot.2016.086

34. Wang J, Zhao Y, Zhou X, Hiebert SW, Liu Q, Shyr Y. Nascent RNA sequencing analysis provides insights into enhancer-mediated gene regulation. BMC Genomics. Aug 23 2018;19(1):633. doi:10.1186/s12864-018-5016-z

35. Chargé SB, Rudnicki MA. Cellular and molecular regulation of muscle regeneration. Physiol Rev. Jan 2004;84(1):209–38. doi:10.1152/physrev.00019.2003

36. Doench JG, Fusi N, Sullender M, et al. Optimized sgRNA design to maximize activity and minimize off-target effects of CRISPR-Cas9. Nat Biotechnol. Feb 2016;34(2):184–191. doi:10.1038/nbt.3437

37. Bassi ZI, Fillmore MC, Miah AH, et al. Modulating PCAF/GCN5 Immune Cell Function through a PROTAC Approach. ACS Chem Biol. Oct 19 2018;13(10):2862–2867. doi:10.1021/acschembio.8b00705

38. Spitz F, Demignon J, Porteu A, et al. Expression of myogenin during embryogenesis is controlled by Six/sine oculis homeoproteins through a conserved MEF3 binding site. Proc Natl Acad Sci U S A. Nov 24 1998;95(24):14220–5. doi:10.1073/pnas.95.24.14220

39. Niro C, Demignon J, Vincent S, et al. Six1 and Six4 gene expression is necessary to activate the fast-type muscle gene program in the mouse primary myotome. Dev Biol. Feb 15 2010;338(2):168–82. doi:10.1016/j.ydbio.2009.11.031

40. Bernecky C, Grob P, Ebmeier CC, Nogales E, Taatjes DJ. Molecular architecture of the human Mediator-RNA polymerase II-TFIIF assembly. PLoS Biol. Mar 2011;9(3):e1000603. doi:10.1371/journal.pbio.1000603

41. Liu Y, Ranish JA, Aebersold R, Hahn S. Yeast nuclear extract contains two major forms of RNA polymerase II mediator complexes. J Biol Chem. Mar 09 2001;276(10):7169–75.

42. Paoletti AC, Parmely TJ, Tomomori-Sato C, et al. Quantitative proteomic analysis of distinct mammalian Mediator complexes using normalized spectral abundance factors. Proc Natl Acad Sci U S A. Dec 12 2006;103(50):18928–33. doi:10.1073/pnas.0606379103

43. Myers LC, Gustafsson CM, Bushnell DA, et al. The Med proteins of yeast and their function through the RNA polymerase II carboxy-terminal domain. Genes Dev. Jan 01 1998;12(1):45–54. doi:10.1101/gad.12.1.45

44. Knuesel MT, Meyer KD, Bernecky C, Taatjes DJ. The human CDK8 subcomplex is a molecular switch that controls Mediator coactivator function. Genes Dev. Feb 15 2009;23(4):439–51. doi:10.1101/gad.1767009

45. Tsai KL, Tomomori-Sato C, Sato S, Conaway RC, Conaway JW, Asturias FJ. Subunit architecture and functional modular rearrangements of the transcriptional mediator complex. Cell. Jun 05 2014;157(6):1430–1444. doi:10.1016/j.cell.2014.05.015

46. Akoulitchev S, Chuikov S, Reinberg D. TFIIH is negatively regulated by cdk8-containing mediator complexes. Nature. Sep 07 2000;407(6800):102–6. doi:10.1038/35024111

47. Lynch CJ, Bernad R, Martínez-Val A, et al. Global hyperactivation of enhancers stabilizes human and mouse naive pluripotency through inhibition of CDK8/19 Mediator kinases. Nat Cell Biol. Oct 2020;22(10):1223–1238. doi:10.1038/s41556-020-0573-1

48. Pelish HE, Liau BB, Nitulescu II, et al. Mediator kinase inhibition further activates super-enhancer-associated genes in AML. Nature. Oct 08 2015;526(7572):273–276. doi:10.1038/nature14904

49. Lynch CJ, Bernad R, Calvo I, Serrano M. Manipulating the Mediator complex to induce naïve pluripotency. Exp Cell Res. Oct 15 2020;395(2):112215. doi:10.1016/j.yexcr.2020.112215

50. Adler AS, McCleland ML, Truong T, et al. CDK8 maintains tumor dedifferentiation and embryonic stem cell pluripotency. Cancer Res. Apr 15 2012;72(8):2129–39. doi:10.1158/0008-5472.CAN-11-3886

51. Lee JC, Liu S, Wang Y, Liang Y, Jablons DM. MK256 is a novel CDK8 inhibitor with potent antitumor activity in AML through downregulation of the STAT pathway. Oncotarget. Nov 02 2022;13:1217–1236. doi:10.18632/oncotarget.28305

52. Laubscher D, Gryder BE, Sunkel BD, et al. BAF complexes drive proliferation and block myogenic differentiation in fusion-positive rhabdomyosarcoma. Nat Commun. Nov 26 2021;12(1):6924. doi:10.1038/s41467-021-27176-w

53. Relaix F, Demignon J, Laclef C, et al. Six homeoproteins directly activate Myod expression in the gene regulatory networks that control early myogenesis. PLoS Genet. Apr 2013;9(4):e1003425. doi:10.1371/journal.pgen.1003425

54. Giordani J, Bajard L, Demignon J, Daubas P, Buckingham M, Maire P. Six proteins regulate the activation of Myf5 expression in embryonic mouse limbs. Proc Natl Acad Sci U S A. Jul 03 2007;104(27):11310–5. doi:10.1073/pnas.0611299104

55. Hsu JY, Danis EP, Nance S, et al. SIX1 reprograms myogenic transcription factors to maintain the rhabdomyosarcoma undifferentiated state. Cell Rep. Feb 01 2022;38(5):110323. doi:10.1016/j.celrep.2022.110323

56. Spassov DS, Atanasova M, Doytchinova I. Inhibitor Trapping in Kinases. Int J Mol Sci. Mar 13 2024;25(6)doi:10.3390/ijms25063249

57. Hopkins TA, Shi Y, Rodriguez LE, et al. Mechanistic Dissection of PARP1 Trapping and the Impact on In Vivo Tolerability and Efficacy of PARP Inhibitors. Mol Cancer Res. Nov 2015;13(11):1465–77. doi:10.1158/1541-7786.MCR-15-0191-T

58. Rohle D, Popovici-Muller J, Palaskas N, et al. An inhibitor of mutant IDH1 delays growth and promotes differentiation of glioma cells. Science. May 03 2013;340(6132):626–30. doi:10.1126/science.1236062

59. Issa GC, Aldoss I, DiPersio J, et al. The menin inhibitor revumenib in KMT2A-rearranged or NPM1-mutant leukaemia. Nature. Mar 2023;615(7954):920–924. doi:10.1038/s41586-023-05812-3

60. Westerling T, Kuuluvainen E, Mäkelä TP. Cdk8 is essential for preimplantation mouse development. Mol Cell Biol. Sep 2007;27(17):6177–82. doi:10.1128/MCB.01302-06

61. Loncle N, Boube M, Joulia L, et al. Distinct roles for Mediator Cdk8 module subunits in Drosophila development. EMBO J. Feb 21 2007;26(4):1045–54. doi:10.1038/sj.emboj.7601566

62. Bruter Alexandra V VEA, Stavskaya Nina I, Antysheva Zoia G, Manskikh Vasily N, Tvorogova Anna V, Korshunova Diana. S, Khamidullina Alvina I, Utkina Marina V, Bogdanov Viktor P, Baikova Iuliia P, Nikiforova Alyona I, Albert Eugene A, Maksimov Denis O, Li Jing, Chen Mengqian, Schools Gary. P, Feoktistov Alexey V, Shtil Alexander A, Roninson Igor B, Mogila Vladislav A, Silaeva Yulia Y, Tatarskiy Victor V. Knockout of cyclin dependent kinases 8 and 19 leads to depletion of cyclin C and suppresses spermatogenesis and male fertility in mice. eLife2024. p. RP96465.

63. Mimoso CA, Goldman SR. PRO-seq: Precise Mapping of Engaged RNA Pol II at Single-Nucleotide Resolution. Curr Protoc. Dec 2023;3(12):e961. doi:10.1002/cpz1.961

64. Skene PJ, Henikoff JG, Henikoff S. Targeted in situ genome-wide profiling with high efficiency for low cell numbers. Nat Protoc. May 2018;13(5):1006–1019. doi:10.1038/nprot.2018.015

