## Supplementary material for "CDK8 Inhibition Releases the Muscle Differentiation Block in Fusion-driven Alveolar Rhabdomyosarcoma": Fig. S

### Figure S1

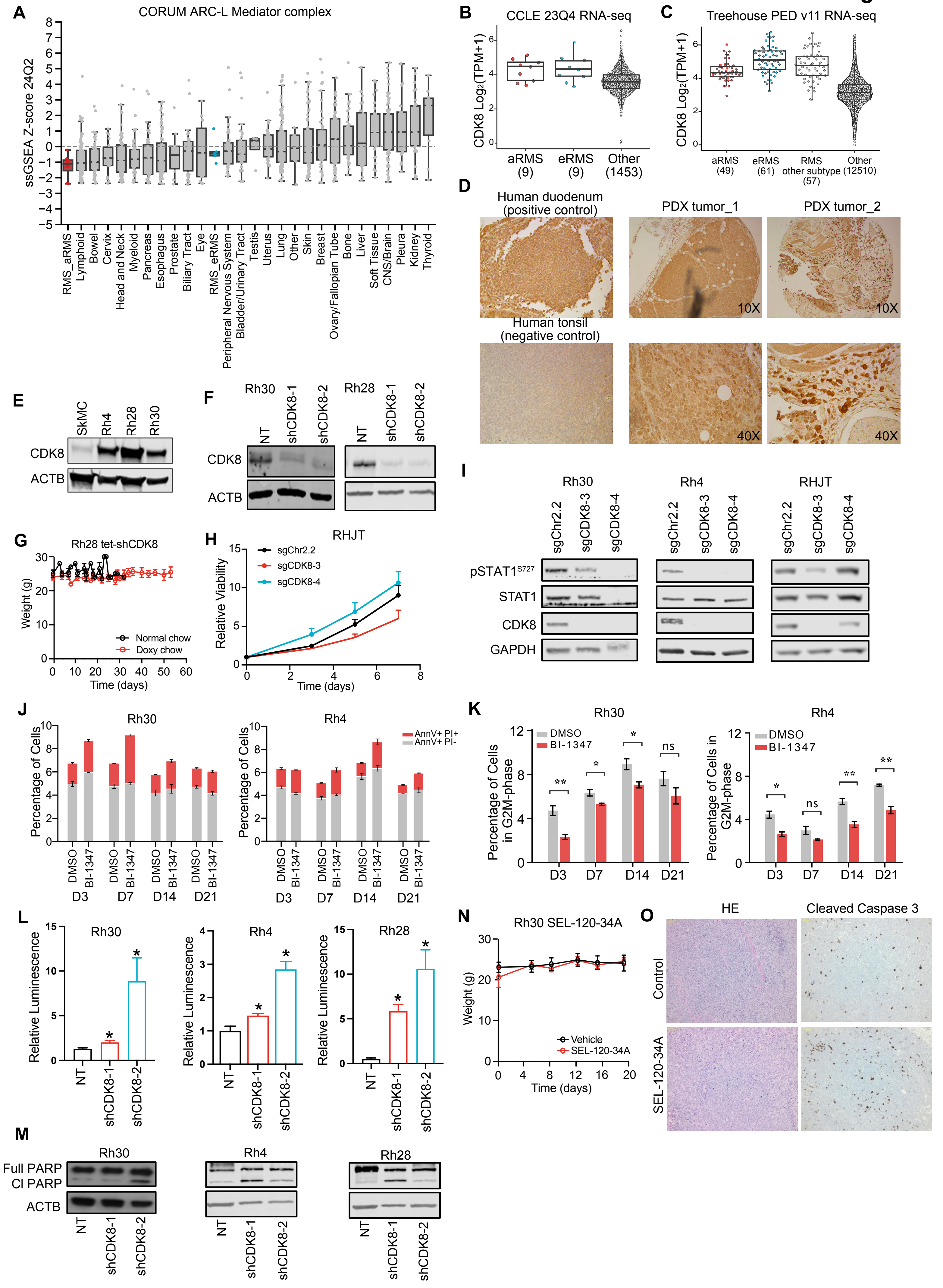
A
CORUM ARC-L Mediator complex
ssGSEA Z-score 24Q2
RMS_aRMS
Lymphoid
Bowel
Cervix
Head and Neck
Myeloid
Pancreas
Esophagus
Prostate
Biliary Tract
Eye
RMS_eRMS
Peripheral Nervous System
Bladder/Urinary Tract
Testis
Uterus
Lung
Other
Skin
Breast
Ovary/Fallopian Tube
Bone
Liver
Soft Tissue
CNS/Brain
Pleura
Kidney
Thyroid
B
CCLE 23Q4 RNA-seq
CDK8 Log2(TPM+1)
aRMS (9)
eRMS (9)
Other (1453)
C
Treehouse PED v11 RNA-seq
CDK8 Log2(TPM+1)
aRMS (49)
eRMS (61)
RMS other subtype (57)
Other (12510)
D
Human duodenum (positive control)
PDX tumor_1
PDX tumor_2
10X
10X
Human tonsil (negative control)
40X
40X
E
SkMC
Rh4
Rh28
Rh30
CDK8
ACTB
F
Rh30
Rh28
NT
shCDK8-1
shCDK8-2
CDK8
ACTB
G
Rh28 tet-shCDK8
Weight (g)
Normal chow
Doxy chow
Time (days)
H
RHJT
sgChr2.2
sgCDK8-3
sgCDK8-4
Relative Viability
Time (days)
I
Rh30
Rh4
RHJT
sgChr2.2
sgCDK8-3
sgCDK8-4
pSTAT1S727
STAT1
CDK8
GAPDH
J
Rh30
Rh4
Percentage of Cells
AnnV+ PI+
AnnV+ PI-
DMSO
BI-1347
D3
D7
D14
D21
K
Rh30
Rh4
DMSO
BI-1347
Percentage of Cells in G2M-phase
D3
D7
D14
D21
ns
L
Rh30
Rh4
Rh28
Relative Luminescence
NT
shCDK8-1
shCDK8-2
N
Rh30 SEL-120-34A
Weight (g)
Vehicle
SEL-120-34A
Time (days)
O
HE
Cleaved Caspase 3
Control
SEL-120-34A
M
Rh30
Rh4
Rh28
Full PARP
Cl PARP
ACTB
NT
shCDK8-1
shCDK8-2

### Figure S2

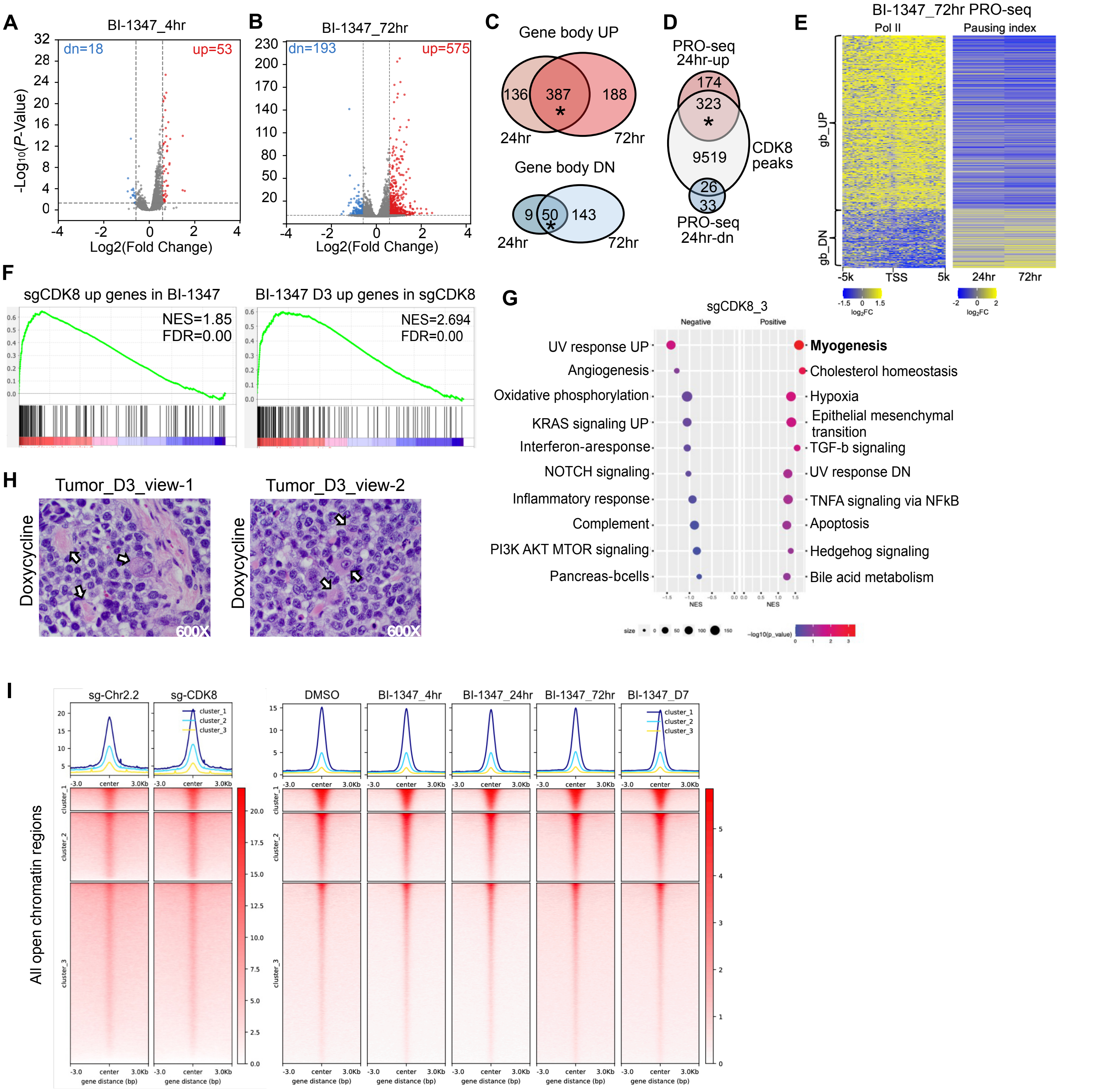
A
BI-1347_4hr
dn=18
up=53
-Log10(P-Value)
Log2(Fold Change)
B
BI-1347_72hr
dn=193
up=575
Log2(Fold Change)
C
Gene body UP
136
387
188
24hr
72hr
Gene body DN
9
50
143
24hr
72hr
D
PRO-seq 24hr-up
174
323
9519
CDK8 peaks
26
33
PRO-seq 24hr-dn
E
BI-1347_72hr PRO-seq
Pol II
Pausing index
gb_UP
gb_DN
-5k
TSS
5k
24hr
72hr
F
sgCDK8 up genes in BI-1347
NES=1.85
FDR=0.00
BI-1347 D3 up genes in sgCDK8
NES=2.694
FDR=0.00
G
sgCDK8_3
Negative
Positive
UV response UP
Angiogenesis
Oxidative phosphorylation
KRAS signaling UP
Interferon-aresponse
NOTCH signaling
Inflammatory response
Complement
PI3K AKT MTOR signaling
Pancreas-bcells
Myogenesis
Cholesterol homeostasis
Hypoxia
Epithelial mesenchymal transition
TGF-b signaling
UV response DN
TNFA signaling via NFkB
Apoptosis
Hedgehog signaling
Bile acid metabolism
NES
H
Tumor_D3_view-1
Tumor_D3_view-2
Doxycycline
600X
I
sg-Chr2.2
sg-CDK8
DMSO
BI-1347_4hr
BI-1347_24hr
BI-1347_72hr
BI-1347_D7
cluster_1
cluster_2
cluster_3
All open chromatin regions
-3.0
center
3.0Kb
gene distance (bp)

### Figure S3

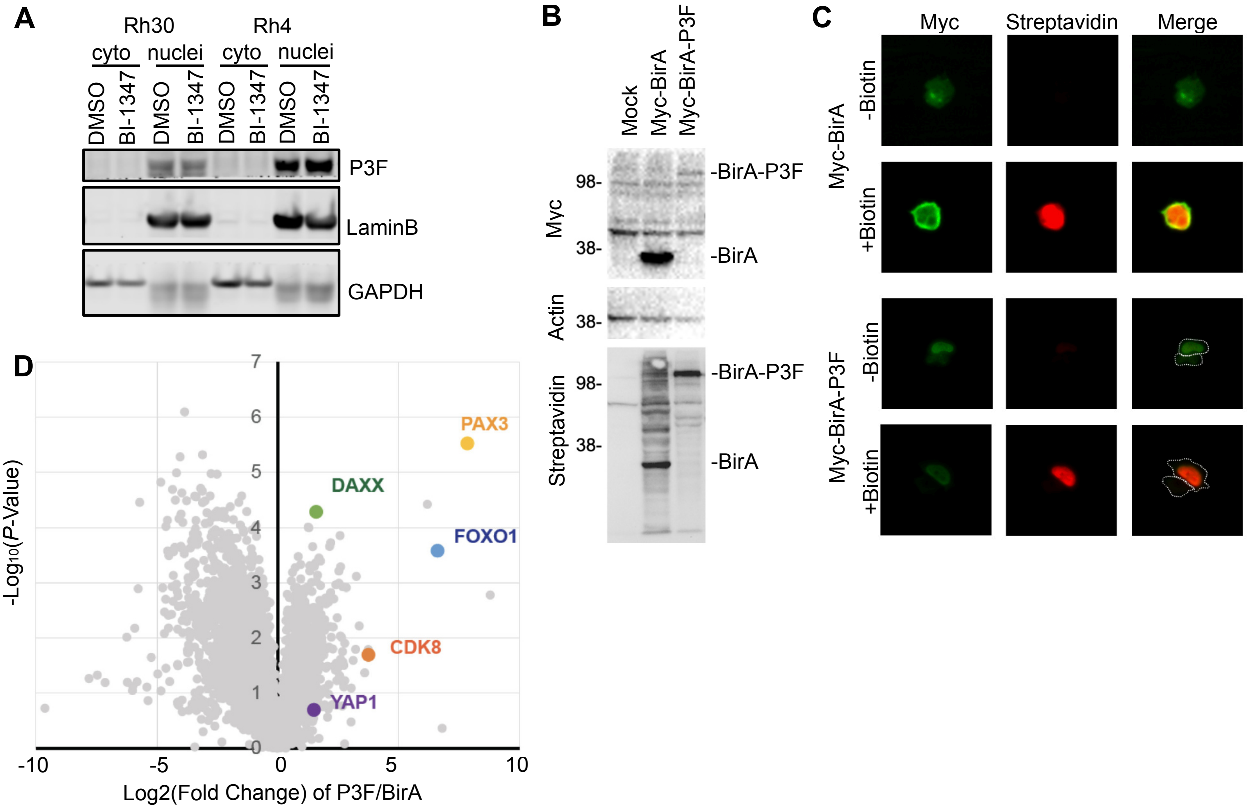
A
Rh30
Rh4
cyto
nuclei
DMSO
BI-1347
P3F
LaminB
GAPDH
B
Mock
Myc-BirA
Myc-BirA-P3F
Myc
Actin
Streptavidin
98-
38-
-BirA-P3F
-BirA
C
Myc
Streptavidin
Merge
Myc-BirA
Myc-BirA-P3F
-Biotin
+Biotin
D
PAX3
DAXX
FOXO1
CDK8
YAP1
-Log10(P-Value)
Log2(Fold Change) of P3F/BirA
-10
-5
0
5
10
1
2
3
4
5
6
7

**Figure S4**

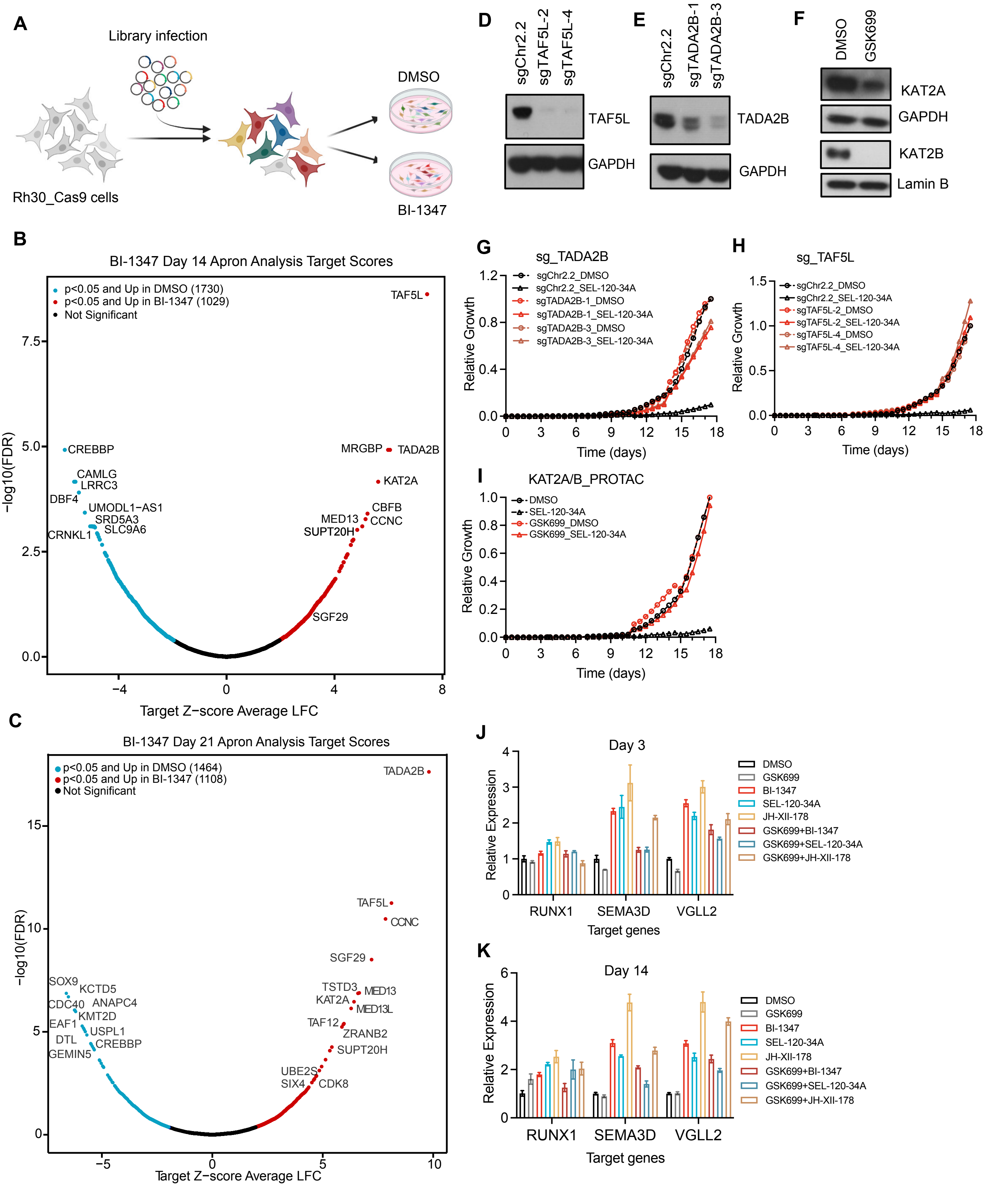
A
Library infection
Rh30_Cas9 cells
DMSO
BI-1347
D
sgChr2.2
sgTAF5L-2
sgTAF5L-4
TAF5L
GAPDH
E
sgChr2.2
sgTADA2B-1
sgTADA2B-3
TADA2B
GAPDH
F
DMSO
GSK699
KAT2A
GAPDH
KAT2B
Lamin B
B
BI-1347 Day 14 Apron Analysis Target Scores
p<0.05 and Up in DMSO (1730)
p<0.05 and Up in BI-1347 (1029)
Not Significant
−log10(FDR)
Target Z−score Average LFC
TAF5L
CREBBP
CAMLG
LRRC3
DBF4
UMODL1−AS1
SRD5A3
SLC9A6
CRNKL1
MRGBP
TADA2B
KAT2A
CBFB
CCNC
MED13
SUPT20H
SGF29
C
BI-1347 Day 21 Apron Analysis Target Scores
p<0.05 and Up in DMSO (1464)
p<0.05 and Up in BI-1347 (1108)
Not Significant
−log10(FDR)
Target Z−score Average LFC
TADA2B
TAF5L
CCNC
SGF29
SOX9
KCTD5
CDC40
ANAPC4
KMT2D
EAF1
USPL1
DTL
CREBBP
GEMIN5
TSTD3
MED13
KAT2A
MED13L
TAF12
ZRANB2
SUPT20H
UBE2S
SIX4
CDK8
G
sg_TADA2B
sgChr2.2_DMSO
sgChr2.2_SEL-120-34A
sgTADA2B-1_DMSO
sgTADA2B-1_SEL-120-34A
sgTADA2B-3_DMSO
sgTADA2B-3_SEL-120-34A
Relative Growth
Time (days)
H
sg_TAF5L
sgChr2.2_DMSO
sgChr2.2_SEL-120-34A
sgTAF5L-2_DMSO
sgTAF5L-2_SEL-120-34A
sgTAF5L-4_DMSO
sgTAF5L-4_SEL-120-34A
Relative Growth
Time (days)
I
KAT2A/B_PROTAC
DMSO
SEL-120-34A
GSK699_DMSO
GSK699_SEL-120-34A
Relative Growth
Time (days)
J
Day 3
Relative Expression
RUNX1
SEMA3D
VGLL2
Target genes
DMSO
GSK699
BI-1347
SEL-120-34A
JH-XII-178
GSK699+BI-1347
GSK699+SEL-120-34A
GSK699+JH-XII-178
K
Day 14
Relative Expression
RUNX1
SEMA3D
VGLL2
Target genes
DMSO
GSK699
BI-1347
SEL-120-34A
JH-XII-178
GSK699+BI-1347
GSK699+SEL-120-34A
GSK699+JH-XII-178

### Figure S5

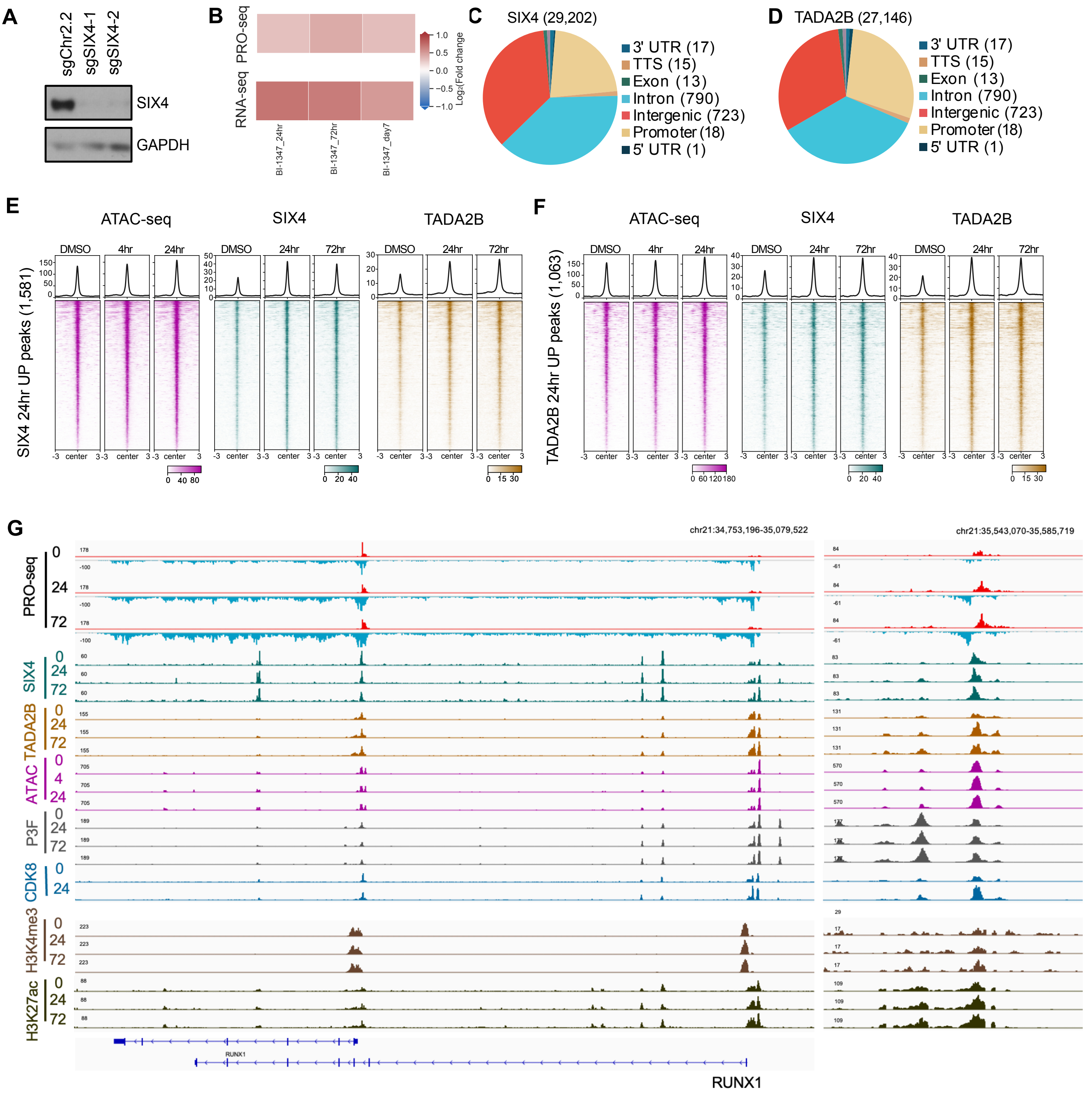
A
sgChr2.2
sgSIX4-1
sgSIX4-2
SIX4
GAPDH
B
PRO-seq
RNA-seq
Log2(Fold change)
BI-1347_24hr
BI-1347_72hr
BI-1347_day7
C
SIX4 (29,202)
3' UTR (17)
TTS (15)
Exon (13)
Intron (790)
Intergenic (723)
Promoter (18)
5' UTR (1)
D
TADA2B (27,146)
3' UTR (17)
TTS (15)
Exon (13)
Intron (790)
Intergenic (723)
Promoter (18)
5' UTR (1)
E
ATAC-seq
SIX4
TADA2B
DMSO
4hr
24hr
72hr
SIX4 24hr UP peaks (1,581)
center
F
TADA2B 24hr UP peaks (1,063)
G
chr21:34,753,196-35,079,522
chr21:35,543,070-35,585,719
PRO-seq
SIX4
TADA2B
ATAC
P3F
CDK8
H3K4me3
H3K27ac
RUNX1

### Figure S6

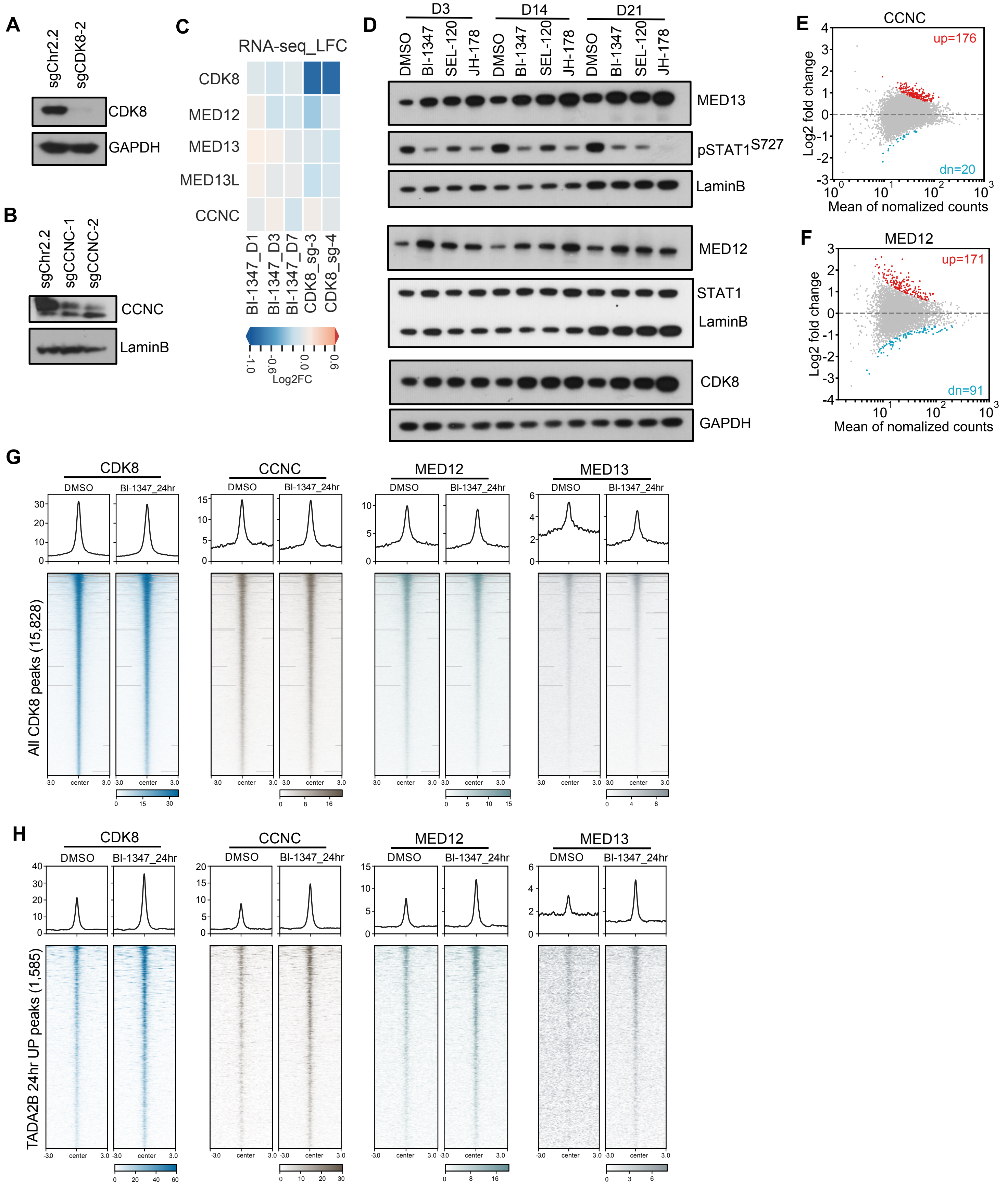

A
sgChr2.2
sgCDK8-2
CDK8
GAPDH
B
sgChr2.2
sgCCNC-1
sgCCNC-2
CCNC
LaminB
C
RNA-seq_LFC
CDK8
MED12
MED13
MED13L
CCNC
BI-1347_D1
BI-1347_D3
BI-1347_D7
CDK8_sg-3
CDK8_sg-4
Log2FC
D
D3
D14
D21
DMSO
BI-1347
SEL-120
JH-178
MED13
pSTAT1S727
LaminB
MED12
STAT1
LaminB
CDK8
GAPDH
E
CCNC
up=176
dn=20
Log2 fold change
Mean of nomalized counts
F
MED12
up=171
dn=91
Log2 fold change
Mean of nomalized counts
G
CDK8
CCNC
MED12
MED13
DMSO
BI-1347_24hr
All CDK8 peaks (15,828)
H
CDK8
CCNC
MED12
MED13
DMSO
BI-1347_24hr
TADA2B 24hr UP peaks (1,585)

#### SUPPLEMENTARY FIGURE LEGENDS

**Figure S1. Genetic or pharmacologic loss of CDK8 triggers minimal apoptosis and cell cycle effects. A.** Boxplots of the ssGSEA scores for CORUM ARC-L-Mediator complex signature comparing aRMS with other cancer cell lines in the Broad Institute DepMap 24Q2 dataset. aRMS is highlighted in red, and eRMS is highlighted in blue. **B, C.** Boxplots showing *CDK8* RNA levels comparing aRMS, eRMS, and other cancer cell lines from the CCLE 23Q4 (**B**) and Pediatric tumors from the Treehouse pediatric cancer (**C**) RNA-seq database. **D.** Human PAX3::FOXO1 fusion-positive aRMS tissue microarrays bearing human PDX tumor were stained for CDK8 using an anti-CDK8 antibody. Human duodenum and tonsil were positive and negative controls, respectively. **E.** Western blot analysis showing the protein levels of CDK8 in aRMS cancer cell lines compared to primary human skeletal muscle cell, SkMC. **F.** Western blot analysis validated the downregulation of CDK8 at the protein level. Actin (ACTB) included as a loading control. **G.** Line graph reveals mean animal weights after inducible knock down of *CDK8* using shRNA. **H.** CRISPR-mediated knockout of *CDK8* by two different gRNAs impairs RHJT aRMS cell growth *in vitro*. Relative growth was assessed by CellTiter-Glo after CRISPR knockout. **I.** Western blot analysis validated decreased CDK8 protein levels and decreased STAT1 serine 727 phosphorylation. GAPDH included as a loading control. **J.** Apoptosis analysis of Rh30 and Rh4 cells treated with DMSO or BI-1347 using flow cytometry analysis for Annexin V and PI staining. Data are presented as mean ± SEM (n=4). **K.** Bar plots showing cell cycle distribution of cells in G2M phase treated with DMSO or BI-1347 for 3, 7, 14, and 21 days using flow cytometry. Data are presented as mean ± SEM (n=4, $p$ derived from an independent T test. *: $p$<=5.0e-02, **: $p$<=1.0e-02, ***: $p$<= 1.0e-03, ****: $p$<=1.0e-04). **L.** Relative caspase activity was measured by Caspase-Glo 3/7 assay after shRNA-mediated *CDK8* perturbation in three indicated aRMS cell lines. Data are presented as mean ± SEM (n=3, $p$ derived from an independent T test. *: $p$<=5.0e-02, **: $p$<=1.0e-02, ***: $p$<= 1.0e-03, ****: $p$<=1.0e-04). **M.** Western blot analysis of full-length and cleaved PARP following *CDK8* knock down by two independent shRNAs or non-targeting control (NT) in Rh30, Rh4, and Rh28 cell lines. ACTB served as a loading control. N. Line graph reveals mean animal weight after treatment of DMSO or SEL-120-34A. **O.** Representative H&E and immunohistochemistry (IHC) staining for cleaved Caspase-3 in tumor sections from control- or SEL-120-34A treated xenograft tumors.

**Figure S2. CDK8 inhibition activates myogenic differentiation-associated gene targets. A. B.** Volcano plots showing the number of gene body changes after 4 hrs (A) or 72 hrs (B) of BI-1347 treatment. Significantly upregulated genes are highlighted in red; significantly downregulated genes are highlighted in blue (padj<0.05, fold change>1.5 or <-1.5). **C.** Venn diagrams showing overlap of genes with up or down regulated gene body changes between the 24- and 72-hour timepoints. **D.** Venn diagrams showing overlap of genes with gene body changes defined by PRO-seq and CDK8 binding. **E.** Heatmaps showing $log_2$ transformed fold change of PRO-seq read counts in 200 bp bins ± 5 kb around the TSS and $log_2$ transformed fold change of the pausing index for all expressed genes at 24- and 72-hour timepoints. **F.** Gene Set Enrichment Analysis (GSEA) comparing gene expression profiles between *CDK8* knockout (sgCDK8) and BI-1347 treatment. **G.** Bubble dot plot of GSEA for gene hits scoring in *CDK8* knockout RNA-seq analysis. **H.** Representative images of H&E stain of xenograft tumors. Arrows indicate myofibrils. **I.** Heatmaps showing the chromatin accessibility of indicated treatment. Heatmaps are clustered by peak intensity (high, medium, and low).

**Figure S3. CDK8 inhibitors induce q differentiation program without altering PAX3::FOXO1. A.** Western blot analysis showing PAX3-FOXO1 protein levels in the cytoplasm or nucleus before or after BI-1347 treatment at indicated time points. Lamin B indicates nuclear localization control, and GAPDH indicates cytoplasmic localization control. **B.** Western Immunoblot of myc-BirA and myc-BirA-PAX3-FOXO1 transiently expressed in HEK293T cells. Streptavidin affinity blot shows unique biotin labeling of myc-BirA-PAX3-FOXO1. **C.** Immunofluorescent images of myc-BirA and myc-BirA-PAX3::FOXO1 in the presence and absence of biotin. White dashed lines represent cell borders. **D.** Volcano plot showing $log_2$ fold change of (myc-BirA-PAX3::FOXO1/Myc-BirA) LC-MS/MS analysis of biotinylated peptides from HEK293T cells expressing myc-BirA and myc-BirA-PAX3::FOXO1.

**Figure S4. Interruption of the SAGA complex rescue the inhibition effect of CDK8 inhibitors. A.** Schematic figure of BI-1347 drug modifier CRISPR screen. **B, C.** Volcano plot of Apron Analysis from genome-scale CRISPR-Cas9 BI-1347 drug modifier screen at day 14 (**B**) and day 21 (**C**). Gene hits highlighted in red indicate sgRNAs significantly enriched in BI-1347 branch, and genes hits highlighted in blue indicate sgRNAs significantly enriched in DMSO branch ($log_2$FC 1.5, padj<0.05). **D-F.** Western blot analysis showing the protein level of TADA2B, TAF5L, and KAT2A/B after CRISPR knock out of *TADA2B* or *TAF5L* or KAT2A/B PROTAC treatment, respectively. GAPDH included as loading control. **G-I.** Live cell proliferation assessed by Incucyte: SEL-120-34A+sgChr2.2, SEL-120-34A+/-sgTADA2B (**G**), SEL-120-34A+/-sgTAF5L (**H**), and SEL-120-34A+/-GSK699 (**I**). **J, K.** Quantitative real-time TaqMan qPCR analysis of *RUNX1*, *SEMA3D*, and *VGLL2* expression at day 3 (**J**) and day 14 (**K**) following treatment with vehicle control (DMSO), CDK8 inhibitors, GSK699, or indicated combination treatments. Expression levels were normalized to *GAPDH* gene expression and shown relative to DMSO control. Data represent means ± SEM (n=6).

**Figure S5. CDK8 inhibition increases SIX4-driven muscle differentiation. A.** Western blot analysis showing SIX4 protein level after *SIX4* knockout by two independent sgRNAs. GAPDH included as a loading control. **B.** Heatmaps of $log_2$ transformed fold change of PRO-seq and RNA-seq read counts of SIX4. **C, D.** Pie charts showing the annotation of all MACS2 defined CUT&RUN peaks of SIX4 (**C**) and TADA2B (**D**) sites after 24 hrs of BI-1347 treatment in Rh30 cells. **E, F.** Heatmaps showing the chromatin accessibility and chromatin occupancy of SIX4 and TADA2B around regions with upregulated SIX4 (**E**) or upregulated TADA2B (**F**) 24 hrs after BI-1347 treatment. **G.** IGV gene tracks showing the PRO-seq, SIX4, TADA2B, ATAC-seq, PAX3::FOXO1, CDK8, H3K4me3, and H3K27ac at the *RUNX1* gene body and its enhancer loci at indicated time points after BI-1347 treatment.

**Figure S6. The Mediator kinase module is required for maximal CDK8 inhibitor activity. A, B.** Western blot analysis showing the protein level of CDK8 (**A**) and CCNC (**B**) after CRISPR knock out, respectively. GAPDH included as loading control. **C.** Heatmap displaying RNA-seq $\log_2$ fold changes in expression of *CDK8* and other Mediator kinase module components (*MED12*, *MED13*, *MED13L*, *CCNC*) in Rh30 cells CDK8 inhibition by BI-1347 or CDK8 knockout by CDK8-targeting sgRNAs. **D.** Western blot analysis showing the protein level of MED13, MED12, and CDK8 after the treatment of three CDK8 inhibitors at indicated time points. GAPDH and Lamin B serve as loading control. **E, G.** MA-plot showing changes of CCNC (**E**) and MED12 (**F**) binding site assessed by CUT&RUN after 24 hrs of BI-1347 treatment. Significantly increased peaks are highlighted in red; significantly decreased peaks are highlighted in blue ($p<0.05$, fold change>1.5 or <-1.5). **G, H.** Heatmaps showing chromatin occupancy of CDK8, CCNC, MED12, and MED13 at all CDK8 binding sites (**G**) or regions with upregulated TADA2B binding (**H**) at 24 hrs of DMSO or BI-1347 treatment.
